# Genome-wide CRISPR screen identifies KEAP1 perturbation as a vulnerability of ARID1A-deficient cells

**DOI:** 10.1101/2023.11.14.566591

**Authors:** LA Fournier, F Kalantari, JP. Wells, JS Lee, G Trigo-Gonzalez, MM Moksa, T Smith, J White, A Shanks, L Wang, E Su, Y Wang, DG Huntsman, M Hirst, PC Stirling

## Abstract

ARID1A is the core DNA binding subunit of the BAF chromatin remodeling complex and is mutated in about ∼8% of all cancers. The frequency of ARID1A loss varies between cancer subtypes, with clear cell ovarian carcinoma (CCOC) presenting the highest incidence at >50% of cases. Despite a growing understanding of the consequences of ARID1A-loss in cancer, there remains limited targeted therapeutic options for ARID1A-deficient cancers. Using a genome-wide CRISPR screening approach, we identify KEAP1 as a genetic dependency of ARID1A in CCOC. Depletion or chemical perturbation of KEAP1 results in selective growth inhibition of ARID1A-KO cell lines and edited primary endometrial epithelial cells. While we confirm that KEAP1-NRF2 signalling is dysregulated in ARID1A-KO cells, we suggest that this synthetic lethality is not due to aberrant NRF2 signalling. Rather, we find that KEAP1 perturbation exacerbates genome instability phenotypes associated with ARID1A-deficiency. Together, our findings identify a potentially novel synthetic lethal interaction of wARID1A-deficient cells.

## INTRODUCTION

AT-rich interaction domain 1A (ARID1A) is the DNA binding subunit of the BAF complex, the canonical SWI/SNF chromatin remodeling complex, which regulates a variety of processes within the cell, including chromatin accessibility, gene expression and the maintenance of genome integrity. ARID1A is mutated in ∼8% of all cancers, with clear cell ovarian carcinoma (CCOC) presenting the highest incidence of ARID1A loss at >50% of cases^1,2^. ARID1A mutations typically occur throughout the length of the gene and result in loss of protein expression. While ARID1A loss has shown prognostic value for several neoplastic malignancies (e.g. gastric^3–5^, lung^6^, hepatocellular^7–9^, and breast cancer^10^), it remains ambiguous how ARID1A status influences the prognosis of gynecologic malignancies. Previous studies have reported adverse clinical outcomes for patients harbouring ARID1A mutations^10,11^, though conflicting reports have emerged suggesting no clinical association (e.g. stage, survival and histopathological features) between ARID1A-positive and negative cohorts^12–17^. Nevertheless, loss of ARID1A is widely recognized as an enabling factor for cancer development and progression.

ARID1A deficiency results in a broad range of phenotypes, including genome instability^18–20^, transcriptional dysregulation^21,22^, and metabolic dysfunction^23–25^. Despite an increasing body of evidence documenting context-specific consequences of ARID1A loss in cancer, there remains a critical gap in knowledge as to how to selectively treat patients harboring these lesions. Genetic dependencies of ARID1A have been used to suggest novel therapeutic avenues against ARID1A-deficient cancer cells. For example, taking advantage of the antagonistic relationship between BAF and the polycomb repressive 2 (PRC2) complex on the regulation of gene expression^26–28^ the EZH2 inhibitor tazemetostat is currently in phase II clinical trials for the treatment of ARID1A-deficient ovarian cancers^29–31^ (EPZ-IST-001). Other promising therapies include BET inhibitors (e.g. JQ1)^32^, agents targeting DNA and replication stress responses (ATR/PARP inhibitors)^18^. The diverse targets of these inhibitors reflect the diversity of cellular activities disrupted by ARID1A loss. Nevertheless, treatment options for CCOC patients harbouring ARID1A mutations remain poorly specific, with surgery and chemotherapy (platinum + taxane) being the standard of care^33,34^.

In this study we employ a genome-wide CRISPR screen to identify genes important for fitness in an isogenic CCOC cell line model of ARID1A loss. We identify the oxidative stress regulator KEAP1 as a potentially novel synthetic lethal partner of ARID1A. Using clinical and cell line samples, we document that KEAP1 perturbation with CRISPR, small molecule inhibitors and RNAi selectively impairs the growth of ARID1A-deficient cells, and results in enhanced genome instability phenotypes. Building on these findings, we show that the combination treatment of ATR inhibitors and small molecules targeting KEAP1 impairs the growth of ARID1A-KO cells, suggesting that the genome instability induced in ARID1A-KO following KEAP1 perturbation can be harnessed therapeutically. We also observed a general dysregulation of KEAP1-NRF2 signalling in ARID1A-KO cells, suggesting that perhaps other functions of KEAP1 underlie the genetic dependency between KEAP1 and ARID1A. Altogether, our findings uncover a novel vulnerability of ARID1A-deficient CCOC cells.

## RESULTS

### Genome-wide CRISPR screen identifies genes important for fitness in ARID1A-KO CCOC cells

To identify genes important for fitness when ARID1A is lost in the context of CCOC, we performed genome-wide CRISPR screens in biological duplicates in RMG-1 cells and its ARID1A knockout (ARID1A-KO) derivative using the Toronto Knockout version 3 library (TKOv3)^35^ (**Figure 1A**). RMG-1 ARID1A-KO cells were generated using CRISPR as documented previously^19^ (**Figure S1A**). Cells were infected at >500X coverage of the library and were cultured until 14 days post puromycin selection to capture robust fitness defects induced by sgRNA knockouts. Genomic DNA was purified and sgRNA representation was assessed by deep amplicon sequencing (>99% sgRNA representation across all samples). Reads were aligned to the TKOv3 library using Bowtie2^36^ and sgRNA enrichment was calculated using the BAGEL2 pipeline^37^, which integrates the fitness scores from all four sgRNAs into a single value for each target gene. Bayes Factor (BF) scores obtained from the BAGEL2 analysis were compared between the ARID1A-WT and ARID1A-KO datasets to identify genes important for fitness, specifically within the ARID1A-KO populations. As expected, non-targeting control sgRNAs targeting eGFP, LacZ and Luciferase presented to lowest BF scores (**Table S1**) across all samples. Furthermore, our approach recovered >94% (ARID1A-WT) and >87.5% (ARID1A-KO) of the training dataset of core essential genes (CEGs) as important for fitness (**Table S1B-C**). As further validation to our experimental approach, precision-recall (PR) curves were plotted to assess screen performance, and the high precision-to-recall ratio suggested a low false positive (precision) and high false-negative rates (recall) (**Figure 1B**).

**Figure 1.**
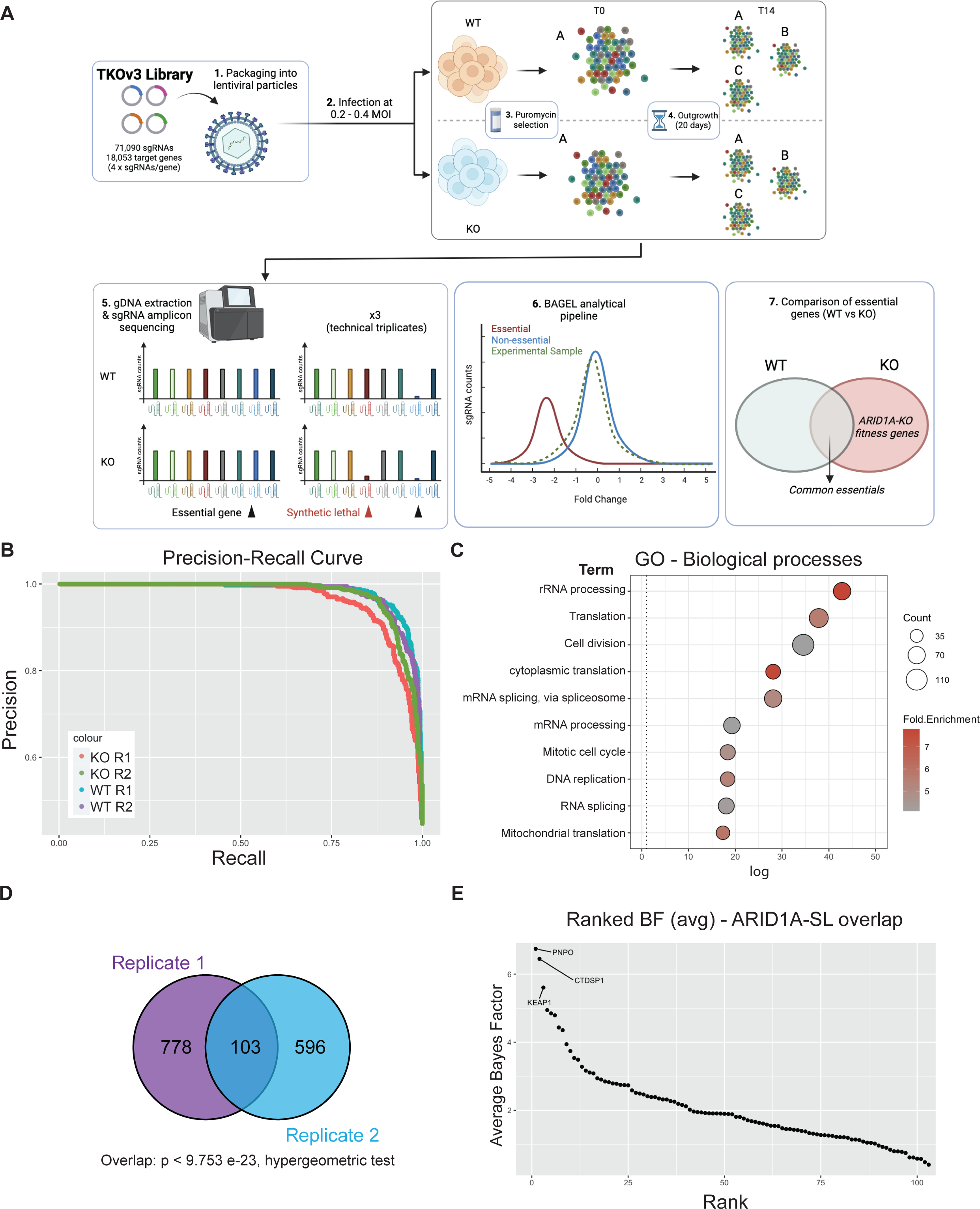
Genome-wide CRISPR screen identifies KEAP1 as a synthetic lethal partner of ARID1A. **(A)** Schematic of CRISPR screening workflow with the TKOv3 library. **(B)** Precision-Recall curve assessing CRISPR screen performance of individual replicates (values obtained from BAGEL2 algorithm, **Supplemental Table 1.5-6**). **(C)** Top 10 enriched Biological Processes GO terms identified using David v 6.8 (p<0.05, with FDR correction) for the ARID1A-KO fitness genes identified by either of our CRISPR screens (n = 1,484 genes, **Supplemental Table 2.1**). **(D)** Overlap of ARID1A-SL hits from both biological replicates of the CRISPR screen. **(E)** Ranked ARID1A-KO specific hits (average BF from both biological replicates, **Supplemental Table 1.8**). The top 3 hits are labeled on the graph.

In total, each biological replicate of our screen identified a consensus of 1,484 genes important for fitness of ARID1A-KO cells (**Supplemental Table 1.7**). To investigate the potential functional repertoire of all the identified hits, we performed a GO analysis on this dataset, which revealed an enrichment in pathways relating to gene expression, cell cycle and mitochondrial function, pathways known to impact fitness in cancer cells ^38–41^ (**Figure 1C and Table S2.1**). To focus on potential genetic dependencies unique to ARID1A-KO cells, we filtered out fitness genes that were also identified in ARID1A-WT cells. This resulted in 881 and 699 ARID1A-KO-specific hits from each replicate, with a significant overlapping consensus of 103 high-confidence synthetic lethal partners, dubbed “ARID1A-SL” (p < 9.753e-23, hypergeometric test, **Figure 1D and Table S1.8** and **S2.2**). Importantly, the overlap of 103 genes successfully identifies known ARID1A synthetic lethal partners in DDX19A^42^, SMARCC1^43^, and FAAP24^42^ amongst others (**Table S1.8**). Additional known synthetic lethal partners were also recovered in either of the screen replicates (e.g. SMARCB1/E1^42,43^, ARID1B^44^, and BRD2^32^; **Table S1.8**).

To prioritize and pursue strong novel candidate synthetic lethal partners of ARID1A for further validation, we averaged the BF score of the ARID1A-SL dataset and ranked the top hits (**Figure 1E**). This approach identified PNPO, CTDSP1 and KEAP1 as the top essential genes when ARID1A is lost. PNPO encodes pyridoxine 5’-phosphate oxidase, which converts dietary vitamin B6 into its biologically active form, contributing to a variety of metabolic processes. CTDSP1 encodes a small phosphatase that targets serine 5 in the heptameric repeats of the C-terminal domain of the large RNA polymerase II subunit POLR2A, controlling gene expression dynamics for most PolII genes. PNPO and CTDSP1 are interesting hits, but likely have pleiotropic effects on global cellularly phenomena. KEAP1 on the other hand is a ubiquitin E3 ligase whose major target, NRF2, a transcription factor with roles in the regulation of metabolism, inflammation, mitochondrial function and more^45^. NRF2’s activity is tightly regulated by KEAP1 under normal conditions, whereby KEAP1 sequesters NRF2 and targets it for proteasomal degradation. Given these data, the fact that ARID1A functionally interacts with NRF2, and that the disruption of this interaction results in fitness defects^24^, we chose to focus on validating KEAP1 as a synthetic lethal partner of ARID1A in CCOC. We further assessed the validity of KEAP1 as a hit by looking at the normalized read counts for all 4 sgRNAs and confirmed the dropout of these guides in the ARID1A-KO samples (**Figure S1D**).

### Validation of KEAP1 as a fitness gene in ARID1A-deficient cells

To validate the potential synthetic lethal relationship between ARID1A and KEAP1, we independently transfected our RMG-1 ARID1A isogenic cell line pair with two independent siRNAs targeting KEAP1 (KEAP1_5 and KEAP1_8), and performed a proliferation assay with crystal violet staining to assess the sensitivity of ARID1A-KO cells to KEAP1 depletion. ARID1A-KO cells showed decreased proliferation following KEAP1 knockdown when compared to their ARID1A-WT parental cells, as measured by the decrease in crystal violet staining (**Figure 2A**). As expected, knockdown of KEAP1 resulted in decreased KEAP1 protein expression that was accompanied by an increase in the levels of its major target protein, NRF2 (**Figure S2A**). Since siKEAP1_5 appeared to yield more consistent knockdowns, we decided to focus on this siRNA moving forward.

**Figure 2.**
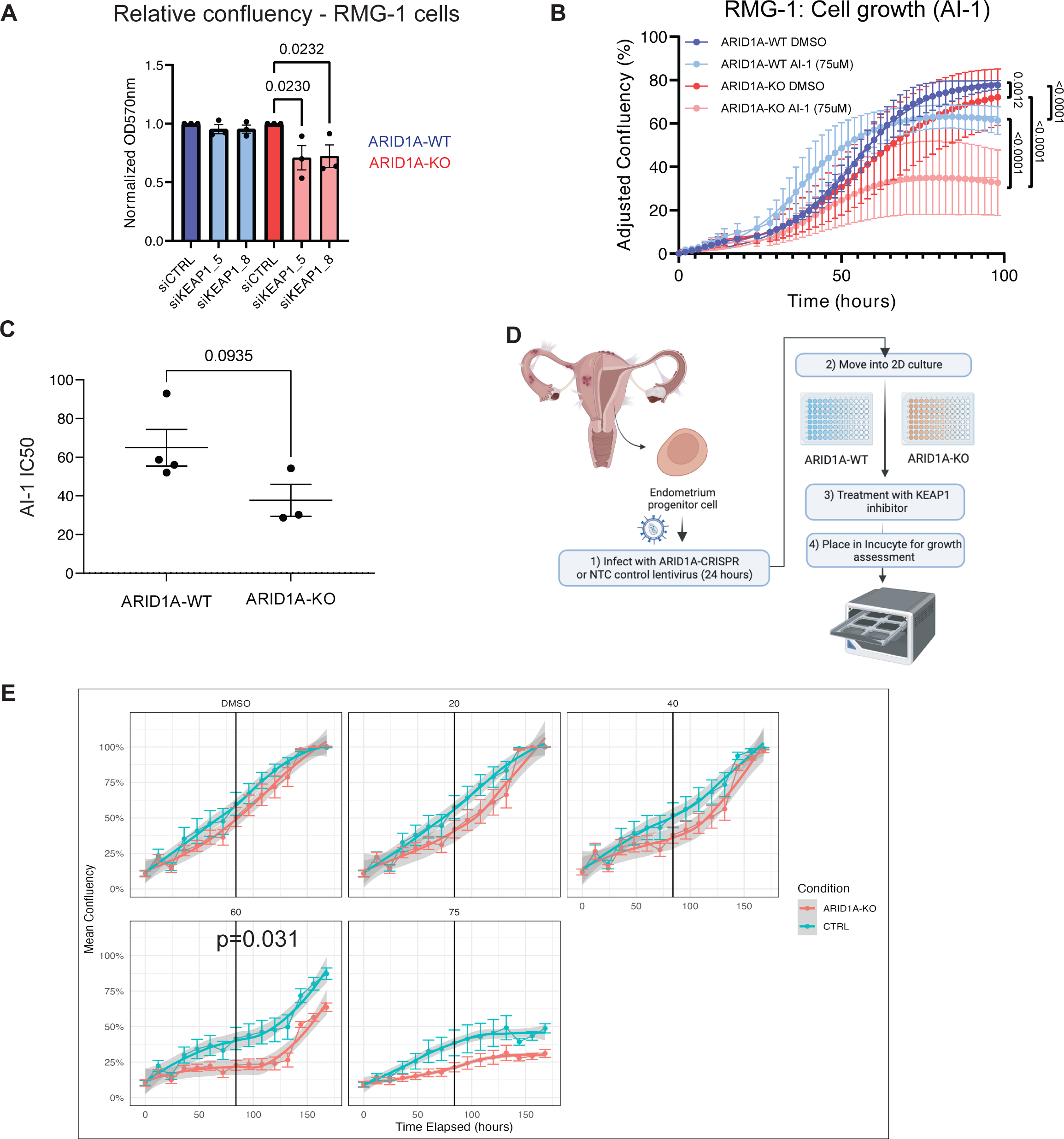
ARID1A-KO cells are sensitive to KEAP1 depletion by siRNA and small molecule perturbation. **(A)** Quantification of crystal violet viability assays in RMG-1 cells (mean ± SEM, one-way ANOVA, n = 3, statistically significant p-values are displayed on graph). **(B)** ARID1A-KO RMG-1 cells grow slower than ARID1A-WT when treated with 75µM AI-1 as measured by IncuCyte S3 imaging system. Relative confluency presented as the growth from initial time point. Error bars represent SEM of averaged triplicated wells from 3 independent experiments. *P* values obtained from extra sum-of-squares F test on calculated logistic growth rate are indicated on graph. **(C)** ARID1A-KO CCOC cells present lower IC50 values to AI-1 than ARID1-WT cells (t-test, mean ± SEM). IC50 values were obtained from dose-response curves presented in **Supplemental Figure S2E** (n = 3). **(D)** Patient-derived endometrium progenitor cell workflow. Normal endometrial tissue was processed and put in 2D culture to expand the progenitor cell population before CRISPR and AI-1 treatments (see Methods). **(E)** Average proliferation curves of three primary endometrial epithelial cell cultures transduced with non-targeting (NTC5) or sgARID1A (ARID1A-KO) lentivirus and treated with AI-1 at the indicated concentrations. Technical quadruplicates from each patient were normalized and combined. p-value was determined at the midpoint for 60µM AI-1 (Welch’s T-test). CRISPR knockdown western blots, and individual patient derived growth curves are shown in **Figure S2E**.

In a parallel approach, we treated ARID1A-deficient or proficient RMG-1 cells with a small molecule targeting KEAP1, called AI-1. AI-1 covalently modifies cysteine residues on KEAP1’s C-terminus to release NRF2 from KEAP1 and disrupt KEAP1’s interaction with the CUL3-RBX1 complex^46,47^. RMG-1 ARID1A-KO cells presented growth defects when subjected to AI-1 treatment when compared to their ARID1A-WT counterparts (**Figure 2B** and **Figure S2B)**. As expected, treatment with AI-1 resulted in an induction of NRF2 protein levels (**Figure S3G**). To further test the potential generalizability of these findings, we assessed the growth of a panel of CCOC cell lines grouped by ARID1A status when subjected to AI-1 treatment and observed a moderate trend where ARID1A-mutated cells presented lower IC50 values compared to the WT counterparts (**Figure 2C** and **Figure S2C-D**).

Additionally, to further assess the ARID1A-KEAP1 synthetic sick/lethal relationship, we obtained primary endometrial epithelial cells and generated ARID1A-KO primary cells using lentiviral CRISPR constructs (**Figure 2D**). Briefly, normal endometrial tissue was processed and put in 2D culture to expand the epithelial cell population. These cells were then transduced with lentivirus containing a CRISPR sgRNA against ARID1A or a non-targeting control (NTC), and ARID1A-expression was validated by western blot (**Figure S2E**). Cell growth of ARID1A-KO and NTC-transduced cells challenged with increasing concentrations of AI-1 confirmed our observations from cell lines, where ARID1A-KO primary cells from three independent donors presented growth defects compared to their NTC counterparts (**Figure 2E** and **Figure S2E**). Altogether these results confirm the observations from our CRISPR screen and suggest that KEAP1 is important for cellular fitness in ARID1A-deficient ovarian/endometrial derived cells.

### Gene expression analysis reveals a dysregulation of NRF2 target genes in ARID1A-KO cells

To gain further insight on the mechanisms underlying the cellular fitness relationship between KEAP1 and ARID1A, we first assessed if and how NRF2 signalling may be disrupted in ARID1A-KO cells. To do so, we revisited RNA-seq data from Wu *et al.*^22^ in RMG-1 cells that either had intact or mutated ARID1A status. Although this isogenic line was generated using different sgRNAs, these cells possess the same genetic background as that used for our CRISPR screens. Differential gene expression (DGE) analysis using DESEQ2^48^ revealed that ARID1A loss caused a significant up-regulation of 531 genes (ARID1A-UP), and down-regulation of 500 genes (ARID1A-DOWN) (**Figure 3A** and **Table S3**). GO analysis using the ENCODE and ChEA datasets on the differentially expressed genes revealed a significant enrichment of NRF2 (also known as NFE2L2) target genes (**Figure 3B**). These results highlight how ARID1A loss results in significant changes in NRF2-mediated gene expression, and perhaps support the notion that KEAP1 signalling is altered in these cells.

**Figure 3.**
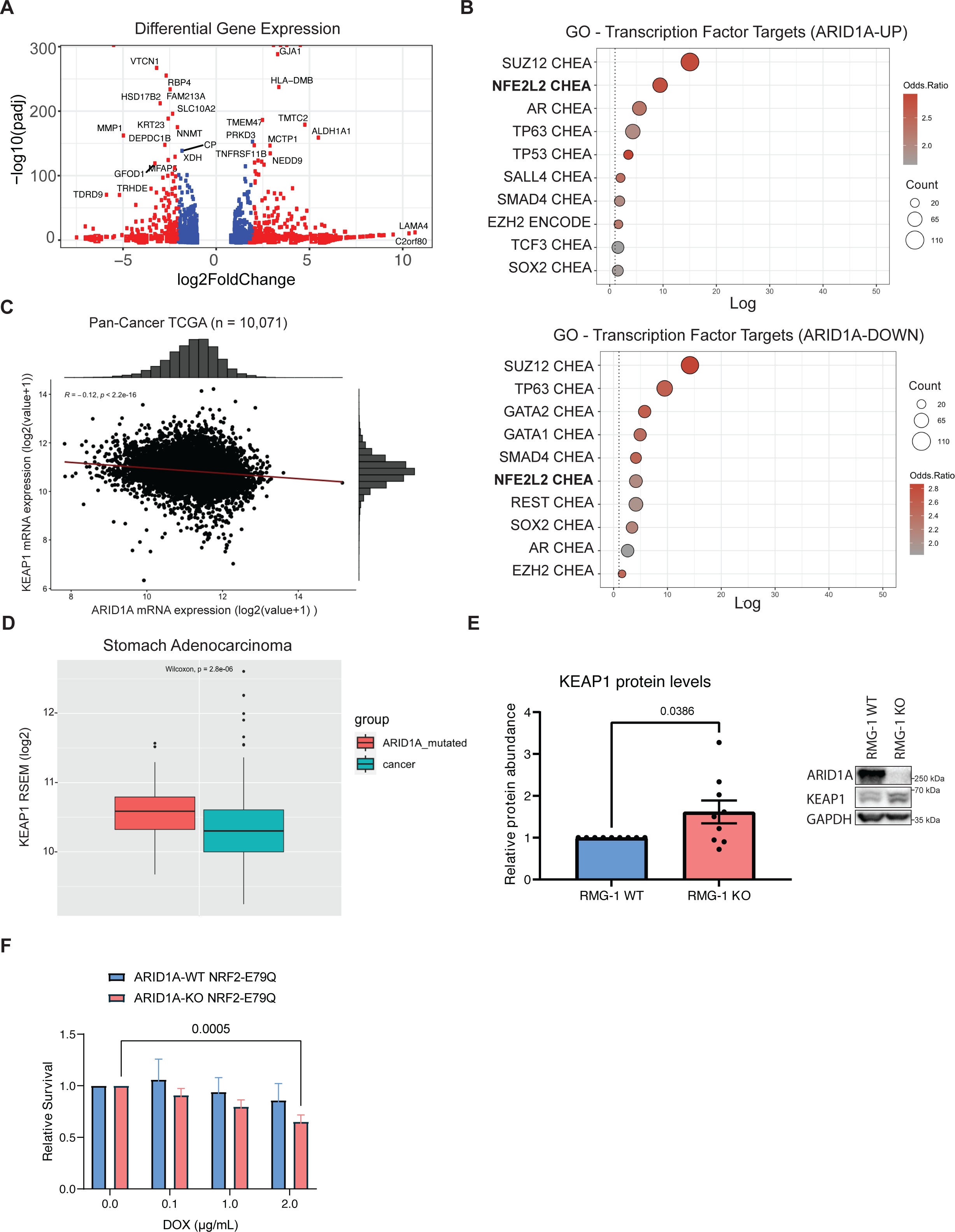
ARID1A-KO sensitivity to KEAP1 depletion is only partly due to dysregulation of NRF2. **(A)** Volcano plot of differentially expressed genes in ARID1A-KO RMG-1 cells from RNA-seq dataset published in Wu *et al.*^22^ (blue dots if padj < 0.01, red dots if log2FC >1 or log2FC < -1 AND padj < 0.01). **(B)** Top 10 enriched transcription factor target genes identified from the ARID1A-UP (*above*) and ARID1A-DOWN (*below*) regulated genes using the ENCODE^76^ and ChEA^77^ datasets (EnrichR^74^; p < 0.05 with Benjamini-Hochberg correction). **(C)** Quantification of KEAP1 and ARID1A mRNA levels (RSEM, log2) from pan-cancer TCGA data (n = 10,071) showing a negative correlation between the two genes. **(D)** Quantification of KEAP1 mRNA levels (RSEM, log2) from TCGA data (stomach adenocarcinoma - stad) showing higher KEAP1 levels in ARID1A-mutated patient samples (n = 90) compared to WT (n = 336) (box limits indicate 25% and 75% over median, Wilcoxon test, p = 2.8e^-6^). **(E)** *Left:* Quantification of KEAP1 protein levels from western blot data showing higher levels in ARID1A-KO RMG-1 cells. KEAP1 intensity was normalized to respective GAPDH loading control, and then to ARID1A-WT (data from 9 from independent samples, mean ± SEM, t test, significant p-values displayed on graph). *Right:* Representative western blot image. **(F)** Quantification of crystal violet viability assay showing that ARID1A-KO RMG-1 cells exhibit sensitivity to NRF2^E79Q^ expression induced by doxycycline (mean ± SEM, ANOVA, n = 3, significant p-values displayed on graph).

Since KEAP1 and ARID1A are commonly mutated in cancer^1,2,49^, we compared mRNA expression of KEAP1 and ARID1A in a large cohort of solid tumours of various origins from The Cancer Genome Atlas (TCGA) datasets (**Figure 3C** and **S3A**). This analysis revealed that transcript levels from KEAP1 inversely correlated with ARID1A, albeit mildly, across a pan-cancer panel (n = 10,071, r = -0.12, p = 2.2e^-16^). Conversely, a weak positive correlation (n = 10,071, R = 0.095, p<2.2e^-16^) was observed between the expression of NRF2 and ARID1A (**Figure S3B**). This observation suggests that ARID1A and KEAP1 could compensate for the loss of each other in cancer development. Furthermore, stratification of patients by ARID1A mutational status confirmed this relationship, where some ARID1A-mutated cancer subtypes (e.g. stomach adenocarcinoma - “stad”) appear to upregulate KEAP1 compared to wild-type (**Figure 3D** and **Figure S3A**). To address this hypothesis, we quantified KEAP1 protein levels from western blots and found a small but significant increase in KEAP1 protein levels in RMG-1 ARID1A-KO cells compared to wild-type (**Figure 3E**). Overall, our data suggests that some ARID1A-deficient cells maintain elevated levels of KEAP1, perhaps as a mechanism to sustain the growth of these cancer cells.

### Aberrant NRF2 signalling does not significantly impair the growth of ARID1A-KO cells

To explore the possibility that aberrant NRF2 signalling may underlie the growth defects of ARID1A-KO cells subjected to KEAP1 perturbation, we generated mutant NRF2-inducible cell lines by transducing our isogenic RMG-1 model (+/- ARID1A) with lentiviral particles containing a pIND20-NRF2^E79Q^-HA construct. The E79Q mutation in NRF2 confers reduced binding affinity for KEAP1, resulting in NRF2 pathway activation^50^. Upon induction of NRF2^E79Q^ protein expression in our ARID1A-WT and KO cells using doxycycline, we observed a minor reduction in the growth of ARID1A-KO cells but not WT cells (**Figure 3F** and **Figure S3C-D**). This phenotype was restricted to the highest dosage of doxycycline (2µg/mL), and we observed no sensitivity to doxycycline at the doses tested (**Figure S3E**). Our data therefore suggests that perhaps the dysregulation of the KEAP1 function beyond NRF2 sequestration, and not NRF2 activation only, may be responsible for the sensitivity displayed by ARID1A-KO cells.

### KEAP1 perturbation exacerbates genome instability in ARID1A-deficient cells

Since the canonical KEAP1 substrate NRF2 does not appear to be responsible for the growth defects observed in ARID1A-KO cells, we sought other explanations. Our group has previously reported that ARID1A-deficient cells experience increased rates of DNA replication stress and DNA damage^19^. To assess whether the sensitivity of ARID1A-KO cells to KEAP1 perturbation could be associated with genome instability phenotypes, we performed immunofluorescence assays to measure p-RPA(ser33) foci formation. p-RPA(ser33) foci formation is a *bona fide* marker of DNA replication stress, and indicates RPA filaments modified by ATR. ARID1A-KO cells presented a higher accumulation of p-RPA(ser33) foci compared to WT cells (**Figure 4A**). Importantly, we observed a significant increase in p-RPA(ser33) foci in ARID1A-KO cells treated with sublethal doses of AI-1 (50µM), but not in ARID1A-WT cells (**Figure 4A**). These results suggest that KEAP1 perturbation may enhance the genome instability phenotypes associated with ARID1A-loss. To assess if the replication stress associated with KEAP1 perturbation also resulted in DNA damage, we performed an immunofluorescence assay probing for γH2AX foci formation. In agreement with our previous results, we observed a significant increase in γH2AX foci formation in our RMG-1 ARID1A-KO cells, and 50µM AI-1 treatment caused a significant increase in γH2AX foci formation specifically in the ARID1A-KO cells, but not the WT (**Figure 4B**).

**Figure 4.**
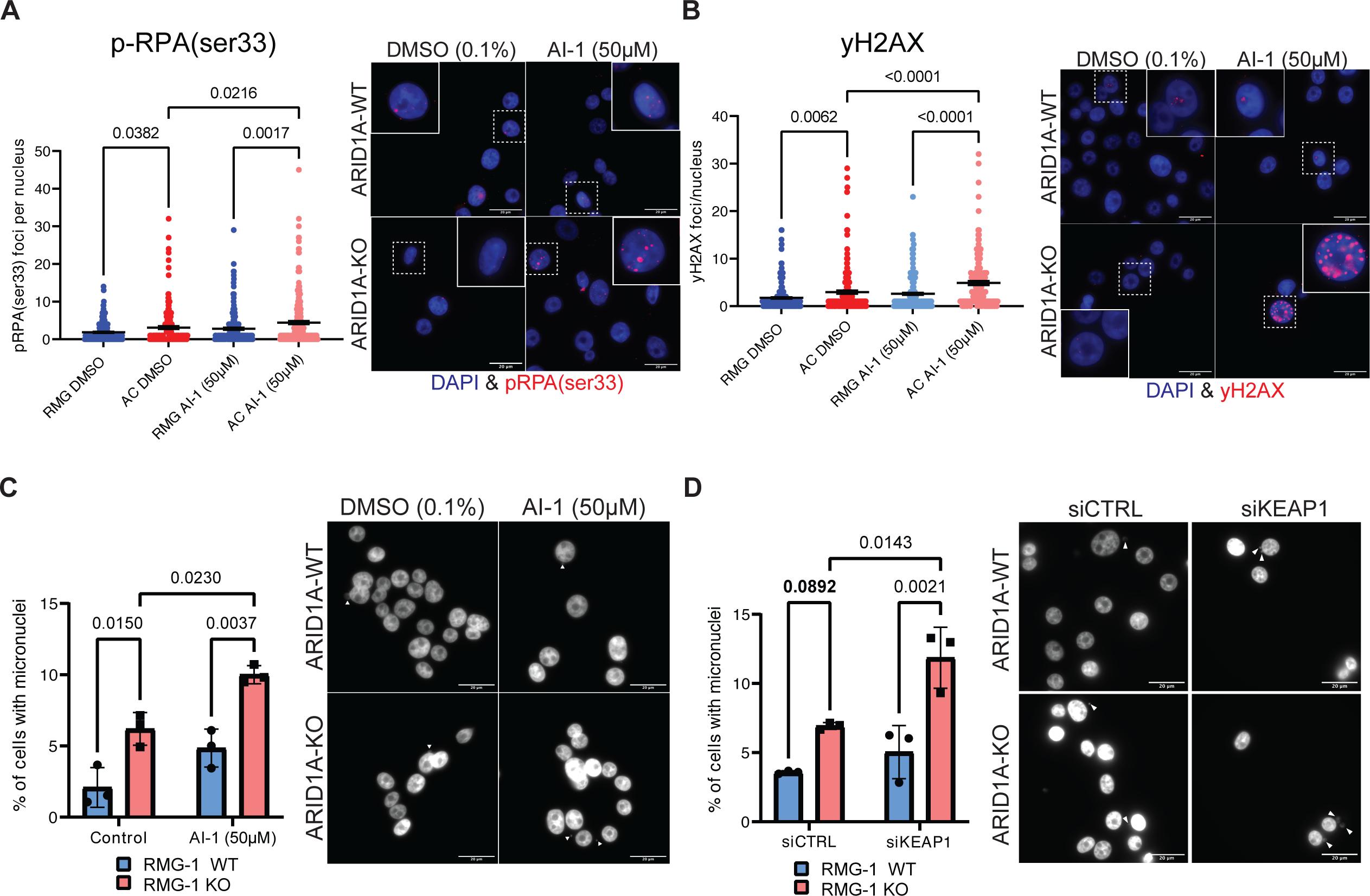
KEAP1 perturbation exacerbates genome instability phenotypes of ARID1A-KO cells. **(A-B)** Quantification (*left*) and representative images (*right*) of immunofluorescence experiment showing AI-1 (50µM) induces higher rates of p-RPAser33 (**A**) or γH2AX (**B**) foci formation in ARID1A-KO RMG-1 cells (mean ± SEM, ANOVA, n = 3, significant p-values displayed on graph). **(C-D)** Quantification (*left*) and representative images (*right*) showing that ARID1A-KO RMG-1 cells accumulate higher levels of micronuclei compared to WT when treated with 50µM AI-1 (C) or siKEAP1 (**D**) (mean ± SEM, ANOVA, n = 3, significant p-values displayed on graph). White arrows highlighting micronuclei.

If DNA damage and replication stress impacts the fitness of ARID1A-KO cells, evidence of induced genome instability should be evident upon KEAP1 depletion. To test this, we assessed micronuclei formation by DAPI staining following KEAP1 perturbation. Micronuclei are typically formed from lagging chromosomes during mitosis or appear in senescent cells due to a defective nuclear envelope^51^. We found that under normal conditions, ARID1A-KO cells present a higher frequency of micronuclei^20^ (**Figure 4C**). AI-1 treatment induced an increase in micronuclei formation specifically in ARID1A-KO cells, while no change could be observed in the ARID1A-WT cells in our isogenic cell line (**Figure 4C**). We observed the same trend in ARID1A-KO RMG-1 cells subjected to siKEAP1, (**Figure 4D**). In addition, western blot analysis probing for γH2AX in a representative primary endometrial progenitor sample treated with NTC- or sgARID1A- and challenged with AI-1 revealed an increase in γH2AX in ARID1A-KO (**Figure S4A**). Thus, we conclude that ARID1A-deficient cells exist in a sensitized state where KEAP1 perturbation with AI-1 or siRNA enhances genome instability phenotypes.

### Dual inhibition of KEAP1 and ATR potentiates killing of ARID1A-KO cells

Since ARID1A-loss sensitizes cells to ATR inhibition^52^, and we observed enhanced marks of replication stress when treating cells with KEAP1 inhibitors, we set out to test whether AI-1 treatment could potentiate the selective killing of these cells. To do so, we treated our isogenic RMG-1 cells with sub-lethal concentrations of ceralasertib, a potent ATR inhibitor, alone or in combination with AI-1 (**Figure S4B**). At the 60 hour time point, neither AI-1 nor ceralasertib treatment resulted in significant killing of RMG-1 ARID1A-KO cells, while the combination treatment of AI-1+ceralasertib resulted in the potentiation of the killing of ARID1A mutants (**Figure S4B**). This data agrees with the genome instability phenotypes induced by KEAP1 perturbation in ARID1A-KO cells and suggests that combination treatment with ATRi could be harnessed to improve the selective growth inhibition of ARID1A-KO cells by AI-1.

## DISCUSSION

Despite a high incidence of mutations across cancers, precision medicine options for patients harboring ARID1A mutations in cancer remain limited and unspecific. To identify genetic dependencies of ARID1A-deficient CCOC cells and to ultimately advise on new synthetic lethal target development, we performed two genome-wide CRISPR screens in an isogenic cell line model of ARID1A loss in RMG-1 cells. Our screen identified known ARID1A synthetic lethal partners in SMARCC1 and DDX19A^42,43^. Known ARID1A synthetic lethal paralogue ARID1B was only identified as significant in one of the two biological replicates, highlighting the noise of this approach and the need for replicates. The overlap between our two screens identified a consensus dataset of 103 genes that are important for fitness when ARID1A is lost (**Figure 1D**). While this overlap was statistically significant, our data also highlights the variability between biological replicates of CRISPR screening experiments.

The top hit from our screen, PNPO is a metabolic enzyme involved in the breakdown of vitamin B6. Despite no reports of ARID1A loss impacting vitamin B6 metabolism, there are many reports highlighting the metabolic consequences of ARID1A mutations^23–25^. It is therefore possible that loss of ARID1A, through metabolic reprogramming, would induce sensitivity to PNPO depletion. Further research will be needed to confirm this hypothesis. The 2nd strongest hit from our screen was assigned to CTDSP1, which functions in the regulation of transcriptional elongation by RNAPII. While we did not test effects of CTDSP1 depletion beyond the screen, there is strong evidence that loss of ARID1A results in defects in transcriptional elongation^21^. In fact, ARID1A deficiency impairs promoter proximal pausing dynamics, a crucial step for productive transcriptional elongation^21^. We hypothesize that inhibition of CTDSP1 would exacerbate this phenotype and perhaps sensitize ARID1A-KO cells to CTDSP1 depletion. While these genes are interesting and warrant further investigation, we decided to focus on the 3^rd^ strongest fitness genes identified by our screen, KEAP1, given that ARID1A physically interacts with its downstream target NRF2, and that the disruption of this interaction was linked to fitness defects^24^.

To validate the findings from our CRISPR screen, we assessed the sensitivity of KEAP1 perturbation by siRNA or by small molecule in an isogenic cell line model of ARID1A loss. AI-1 was initially characterized as an NRF2 activator, acting by covalently modifying cysteine residues of KEAP1, to promote the stabilization and transcriptional activation of NRF2^46^. AI-1 also disrupts the ability of KEAP1 to serve as an adaptor for CUL3-KEAP1 ubiquitin ligase complex, potentially impacting other functions of KEAP1 like the regulation of protein homeostasis^46^. KEAP1 perturbation by AI-1 resulted in growth defects in a cell line model when ARID1A is lost, and also in primary endometrial progenitors depleted for ARID1A. This was further corroborated in a panel of CCOC cell lines, where we saw a moderate correlation between lower AI-1 IC50 values and ARID1A-deficiency. Overall, our findings suggest that KEAP1 perturbation impairs the growth of ARID1A-deficient cells.

To gain further insight on the potential mechanism underlying the sensitivity of ARID1A-KO cells to KEAP1 perturbation, we investigated whether NRF2 signalling, downstream KEAP1, could explain the growth defects we observed. Building on work by Wu *et al.*^22^, we performed a differential gene expression analysis that identified large subsets of genes that are disrupted when ARID1A is lost. Among these, polycomb repressive complex 2 (PRC2) target genes were highly enriched within the DGE dataset, as seen by the enrichment of the PRC2 subunits SUZ12 and EZH2 (**Figure 3B)**. This is unsurprising considering that PRC2 and the BAF complex have antagonistic roles in the regulation of gene expression^26,55^. Accordingly, EZH2 inhibition in ARID1A-deficient cells has been reported to behave in a synthetic lethal manner^56,57^. Interestingly, our DGE analysis also revealed that NRF2 (NFE2L2) target genes are highly enriched among both the up-regulated, and to a lesser extent, down-regulated genes (**Figure 3B**). Furthermore, the analysis of pan-cancer TCGA data revealed a mild, but significant correlation where some ARID1A-mutated tumour subtypes present elevated KEAP1 mRNA levels (e.g. stomach adenocarcinoma – STAD, and colon adenocarcinoma – COADREAD) (**Figure 3D** and **Figure S3A**). Unfortunately, limitations in sample size and data availability prevented us from assessing this trend in ovarian carcinoma subtypes specifically. These results suggest that KEAP1 is perhaps up-regulated when ARID1A is mutated, and this may support cancer cell growth.

Although we cannot rule out that transcriptional rewiring by NRF2 activation may be partially responsible for the growth defects of ARID1A-KO cells following KEAP1 perturbation, our findings suggest that functions of KEAP1 beyond NRF2 sequestration may underlie the genetic dependency we report here. To this extent, we demonstrated that KEAP1 perturbation enhances genome instability phenotypes in ARID1A-KO cells, though the nature of the damage sustained by these cells remains unknown. Marzio *et al.*^58^ have shown that loss of KEAP1 can produce a BRCA-ness phenotype, which may explain our observations of elevated genome instability phenotypes in ARID1A-KO cells subjected to KEAP1 perturbation. Moreover, KEAP1 has been shown to interact with proteins like BRCA1 and PALB2, which may influence DNA repair pathway choice and cell cycle regulation. It is possible that defects in DNA repair pathway choice may also result in inadequate DNA repair, though more research is necessary to confirm this hypothesis. Furthermore, two groups have identified MCM3, a component of the replicative DNA helicase complex, as a substrate of KEAP1^59,60^. While the exact function of this interaction remains unknown, Mulvaney *et al*. showed that KEAP1 associates with chromatin during S-phase. The authors suggest that KEAP1 plays a role in monitoring replication fork progression and the coordination of replisome activity. Overall, these results point towards an NRF2-independent role for KEAP1 in the maintenance of genome integrity, which may be further exacerbated when ARID1A is lost. Finally, since KEAP1 is an essential component of the KEAP1-CUL3-RBX1 ubiquitin ligase complex that targets various proteins for proteasomal degradation, and also interacts with p62 to regulate autophagic flux^61–65^, it also remains possible that KEAP1 perturbation could disrupt normal protein homeostasis, indirectly impairing the growth of ARID1A-KO cells.

While our group is the first to our knowledge to document a genetic dependency between KEAP1 and ARID1A, consequences of ARID1A loss on NRF2 signalling have been reported. Ogiwara *et al.*^24^ have reported that ARID1A-loss causes glutathione metabolism deficiencies, which sensitizes ARID1A-KO cells to SLC7A11 depletion. The authors showed that this synthetic lethal relationship was partially explained by the loss of an interaction between ARID1A and NRF2, which in turn regulates SLC7A11 expression. While the role of KEAP1 in this genetic dependency has not been explored, additional evidence suggests that KEAP1-mediated signalling is important in ARID1A-deficient cells. In fact, bromodomain and extra-terminal domain inhibitors (BETi) have been used to selectively kill ARID1A-KO cells; particularly the inhibitor JQ-1, which targets BRD4^32^. While the authors suggest that this toxicity is due to transcriptional changes associated with an increased BRD4 recruitment to chromatin, BRD4 inhibition by JQ-1 has been shown to affect KEAP1-NRF2 signalling, supporting the idea that KEAP1 is perhaps a regulator of fitness in ARID1A-deficient cells^66–68^. Additionally, loss of other BAF complex subunits has been associated with the dysregulation of KEAP1-NRF2 signalling. For instance, mutations in the catalytic subunit BRG1 have been associated with changes in NRF2-mediated gene expression^69^. Overall, this suggests that there exists a convergence in functions between the BAF complex and KEAP1-NRF2 signalling, though further research is needed to explore the nature of this connection. Overall, our we propose a model (**Figure 5**) where KEAP1 perturbation results in an upregulation of NRF2 signalling and perhaps dysregulation of other cellular processes such as protein homeostasis, which is viable when ARID1A is intact; however, when KEAP1 is perturbed in ARID1A-deficient cells, the resulting genome instability is exacerbated by KEAP1 perturbation and results in lethality.

**Figure 5.**
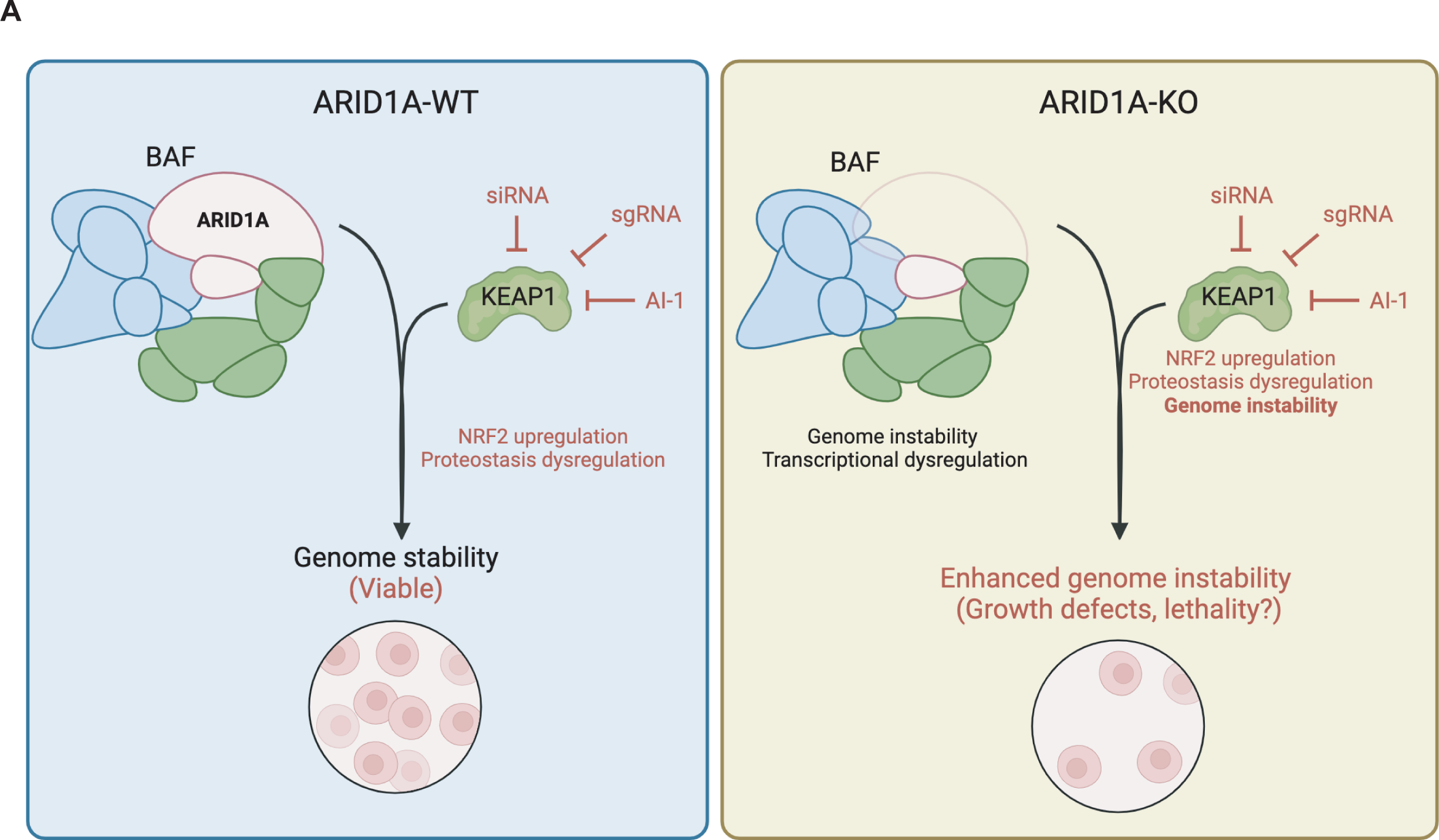
Sensitivity of ARID1A-KO cells to KEAP1 perturbation is conserved in endometrium progenitor-derived organoid model. **A)** Model of ARID1A-KEAP1 synthetic lethality. Perturbation of KEAP1 dysregulates normal NRF2 signalling and proteostasis, which remains viable in normal cells. Perturbation of KEAP1 in ARID1A-KO cells specifically enhances genome instability phenotypes, resulting in growth defects.

We recognize that relatively high concentrations (micromolar range) of AI-1 are required to elicit the phenotypes observed, suggesting that this inhibitor is not promising for clinical use. In fact, the detailed consequences of AI-1 inhibition remain poorly characterized. While some off-target effects have been predicted against HDAC1^46^, our group did not observe significant changes in protein expression when subjecting cells to lethal doses of this small molecule, though these observations do not exclude the possibility that HDAC1 function may be affected by this inhibitor (**Figure S3F**). Nevertheless, our observations suggest that further investigation of KEAP1 function in ARID1A-deficient cells is warranted to further inform on the fitness defects described herein. In fact, our observations that sublethal concentrations of AI-1 in combination with ceralasertib allowed to selectively kill ARID1A-KO cells suggests that combination therapy may potentiate the effects of AI-1, allowing perhaps limit off target effects. Importantly, the validation of our findings using siRNA knockdown of KEAP1 as a complementary approach, supports our conclusions. Ultimately, both generating more potent KEAP1 inhibitors and understanding the mechanisms underlying the growth defects of ARID1A-KO cells is needed to confirm the clinical potential of these findings.

## METHODS

### Cell culture conditions

RMG-1 ARID1A-WT and ARID1A-KO cells were a generous gift from the Huntsman Lab. The CCOC cell line panel (JHOC-5/7, OVISE, OVMANA, OVCA 429 and RMG-II) were also obtained from the Huntsman Lab. All CCOC cell lines were cultured in RPMI 1640 supplemented with 10% FBS and 1% penicillin/streptomycin at 37°C, supplied with 5% CO_2_.

### Generation of NRF2^E79Q^ dox-inducible cell lines

The pIND20-NRF2^E79Q^-HA plasmid was a generous gift from Bernard Weissman. The plasmid was transformed into bacteria for amplification and validated by sequencing (**Table S4, Sheet 2**). To produce lentiviral particles for stable expression of the construct, 8x10^6^ HEK-293T cells were transfected in a 15-cm culture dish. Cells were transfected using the TransIT-LT1 Transfection reagent (Mirus, MIR 2305, 3µL/µg of DNA) with 4.8μg psPAX2, 3.8μg pMD2.G and 8µg pIND20-NRF2^E79Q^-HA DNA 24-hours after cells were seeded. After 24 hours of transfection, the media was replaced with high-BSA growth media (DMEM + 1.1g/100mL BSA + 1X Pen/Strep) for viral harvest. Lentiviral particles were harvested 48-hours post-transfection, and viral particles were concentrated by ultracentrifugation (25,000g, 4°C, 1.5 hours), resuspended in OPTI-MEM and frozen at -80°C for future use. To generate the stable cell lines, 1x10^6^ RMG-1 cells were infected at a MOI between 0.2-0.4 in the presence of 8µg/mL polybrene. The infected cells were selected for 96 hours in media containing 800 µg/mL geneticin (Thermo CAT: 10131027).

### TKOv3 library cloning and viral preparation

The Toronto human knockout pooled library (TKOv3) was a gift from Jason Moffat (Addgene #90294). The library was amplified in bacteria as described in the Moffat protocol on Addgene (https://www.addgene.org/pooled-library/moffat-crispr-knockout-tkov3/). The amplified sgRNA library was packaged into lentiviral particles using HEK293T/17 cells by co-transfection with psPAX2 and pMD2.G viral plasmids (1:1:1 molar ratio). To produce viral particles at scale, 510x10^6^ cells were transfected in 60 15-cm culture dishes. Each dish was transfected using the TransIT-LT1 Transfection reagent (Mirus, MIR 2305, 3µL/µg of DNA) with 4.8μg psPAX2, 3.8μg pMD2.G and 8μg library DNA 24-hours after cells were seeded. After 24 hours of transfection, the media was replaced with high-BSA growth media (DMEM + 1.1g/100mL BSA + 1X Pen/Strep) for viral harvest. Lentiviral particles were harvested twice at 48-hours and 72-hours post-transfection. Both harvests were pooled, viral particles were concentrated by ultracentrifugation (25,000g, 4°C, 1.5 hours) and frozen at -80°C for future use.

### TKOv3 library viral transduction and titration

Viral titre was assessed to determine the multiplicity of infection (MOI) of the concentrated viral particles. RMG-1 ARID1A-WT and ARID1A-KO cells were seeded at a density of 2.5x10^6^ cells/well in 12-well plates and spin-fected (2000 rpm for 2 hours at 37°C) with increasing concentrations of virus in the presence of 8µg/mL polybrene (Millipore TR-003-G lot#3287963). The next day, the cells were split into 6-well plates at a density of 0.5x106 cells/well and subjected to 2µg/mL puromycin (Sigma P8833) for selection of infected cells. Cells not transduced with the library (no virus control) did not survive past 24 hours of puromycin selection (2µg/mL). After 48-hours of selection, all cells in all wells were counted to determine the viral volume that resulted in 20-40% survival in puromycin (corresponding to an MOI of 0.2-0.4 assuming an independent infection rate).

### CRISPR knockout pool generation

The dropout CRISPR screen was performed in biological duplicates, in an adapted version of the Mair *et al.* protocol^72^. ARID1A-WT and ARID1A-KO RMG-1 cells were transduced with sgRNA libraries at a multiplicity of infection of (MOI) between 0.2 and 0.4, aiming for coverage of, on average, >500 cells per sgRNA reagent. For genome-wide screens, 120 million cells were transduced by “spin-fection” (2 hours, 2000rpm, 37°C) per cell line in 12-well plates (2.5x10^6^ cells per well) using the appropriate volume of viral particles for MOI of 0.2-0.4 and 8µg/mL polybrene. Two additional wells per cell line were also plated to account for no puromycin and no virus controls to assess MOI at time 0 (T0). Media was aspirated and refreshed following the infection and cells were incubated overnight. The following, the cells were split, pooled, and seeded at 8x10^6^/plate in 150mm dishes in media containing 2µg/mL puromycin. Additional plates were seeded to assess MOI (no-puromycin, no-virus and virus-infected). Forty-eight hours post-selection with puromycin, the cells were split, and MOI was assessed. Time 0 samples were frozen to assess initial library representation (20x10^6^ cells - >250X), and the remainder of cells were maintained in culture and split as needed to ensure confluence did not exceed 90%. Cells were harvested and frozen as “dry” pellets (-80°C) at relevant time points for subsequent sgRNA enrichment assessment (described below).

### Genomic DNA extraction and precipitation

Genomic DNA extraction and precipitation was performed according to the method described in the TKOv3 protocol described by Mair *et al.*^72^ using the QIAamp Blood Maxi kit (cat no. 51194) and RNase A (cat no. 19101).

### Library preparation and sequencing

The sgRNA libraries were prepared using two-steps of PCR as described by Mair *et al.*^72^: 1) amplify the sgRNA region within genomic DNA; 2) amplify sgRNAs with Illumina TruSeq adapters with i5 and i7 indices (see **Table S4**). These indices are unique sequences that are added to DNA samples during library preparation and act as sample identifiers during multiplex sequencing. All PCR steps were performed using the NEBNext Ultra II Q5 Master Mix high-fidelity polymerase.

For PCR #1, the thermocycling parameters were 98°C for 30s, 25 cycles of (98°C for 10s, 66°C for 30s, 72°C for 15s), and 72°C for 2 minutes. In each PCR #1 reaction, we used 3.5μg of gDNA. For each sample, the necessary number of PCR #1 reactions was used to capture the appropriate representation of the screen. For instance, assuming a diploid genome is ∼7.2 pg and one guide-RNA per genome, 100 µg of genomic DNA yields ∼200-fold coverage of the TKOv3 library. In this example, at least 29 PCR#1 reactions were necessary to capture 200X representation of the library for each replicate.

PCR #1 products were pooled for each biological samples and 5 µL was used for amplification and unique barcoding in PCR #2. The thermocycling parameters for PCR#2 were 98°C for 30s, followed by 10 cycles of (98°C for 10s, 55°C for 30s and 65°C for 15s), and 65°C for 5 minutes. The PCR #2 products were validated using gel electrophoresis and purified using the QIAquick Gel extraction kit (QIAGEN). The purified libraries were then quantified by Nanodrop and Qubit prior to sequencing on an Illumina MiSeq (2 samples per lane for >100X coverage).

### Amplicon scoring, guide-RNA enrichment and hit identification

Deep amplicon sequencing data was processed for sgRNA representation using custom scripts. To summarize, the sequencing reads were first de-multiplexed using the 8-bp barcodes in the forward primer, then using the 8-bp barcodes in the reverse primers. De-multiplexed reads were trimmed to leave only the 20-bp spacer (sgRNA) sequences. The sgRNA sequences were then mapped to the reference TKOv3 library sgRNA sequences using Bowtie 2^48^. For mapping, no mismatches were allowed, and mapped sgRNA sequences were quantified by counting the total number of reads.

Fitness scores were assigned to each gene by integrating the read counts from individual sgRNAs using BAGEL2 (each replicate analysed independently)^37^. Genes with a positive Bayes Factor and an FDR cut-off of 0.05 were determined important for fitness across datasets, and hits that were unique to the ARID1A-KO datasets (i.e. not overlapping between ARID1A-WT and KO) were identified as synthetic lethal to ARID1A. The consensus dataset of synthetic lethal partners of ARID1A corresponds to the intersection of ARID1A-KO specific hits from both biological replicates of the screen.

### Gene ontology (GO) analysis

Gene ontology analysis for biological processes was performed using DAVID v6.8^73^. Gene ontology analysis for target genes of transcription factors was performed using the ENCODE and ChEA datasets from EnrichR^74^.

### Incucyte growth assay

Cells were seeded at 10,000 (RMG-1) cells per well in a clear bottom 96-well plate on day 0. For drug treatments, cells were treated with indicated drug concentrations on day 1 and placed in an IncuCyte S3 live-cell imaging system contained in an incubator kept at 37 °C and 5% CO2. Images were taken at 2-hour intervals in quadruplicate/well for 72-96 hours.

### RNA interference

2 × 10^5^ cells were reverse-transfected in 6-well plates with Flexitube siRNA constructs (Qiagen) targeting KEAP1 (KEAP1_5 or KEAP1_8), NRF2 (NRF2_10), or a non-targeting control (siCTRL) at concentrations of 25 nM with HiPerFect Transfection Reagent (Qiagen, #301705) according to the manufacturer’s protocol. Target sequences are available in **Table S4**. Cells were cultured for 48 h after transfection and before subsequent analysis. Alternatively, for reverse transfection, cells were seeded at 10,000 (RMG-1) cells per well in 24-well plates on day 0. On day 1, cells were transfected with 25nM siRNA (final concentration).

### Crystal violet proliferation assay

*KEAP1 siRNA proliferation:* For siRNA experiments, cells were seeded at 10,000 (RMG-1) cells per well in 24-well plates on day 0. On day 1, cells were transfected with 25nM siRNA (final concentration). Cells were fixed for crystal violet staining after 7 days of outgrowth on Day 8.

For experiments with drug treatments, cells were seeded at 5,000 (OVCA 429) or 10,000 (RMG-1, RMG-II, OVMANA, OVISE, JHOC-5/7) cells per well in 24-well plates, and incubated overnight prior to addition of the drug. Cells were fixed for crystal violet staining after outgrowth of 7 days on day 8.

For experiments with RMG-1 p.IND20-NRF2-E79-HA expressing cells, 10,000 cells were seeded in a 6-well plate on Day 0. On day 1, doxycycline was added to the media at concentrations indicated in Figure legends. Media was changed on day 4 to replenish doxycycline. Cells were fixed for crystal violet staining on day 7.

Briefly, cells were washed with PBS and fixed with 3.7% formaldehyde for 15 minutes at RT. Cells were washed with PBS and stained with crystal violet for 30 minutes at RT, after which cells were washed with distilled H_2_O until no residual crystal violet could be observed. The plates were dried at RT overnight. The following day, the plates were incubated for 15 minutes at RT (in dark) following the addition of 500µL of 10% acetic acid in methanol. The absorbance at 570nm was measured using a Tecan Infinite M200Pro plate reader.

### Immunofluorescence

For all immunofluorescence experiments, cells were grown on coverslips overnight. For experiments with drug treatments, cells were treated at indicated concentrations for 24 hours prior to fixation. For siRNA transfections, cells were reverse transfected at time of seeding with appropriate siRNA concentrations and grown for 48 hours prior to fixation.

Cells were fixed with 4% paraformaldehyde for 10 minutes and permeabilized with 0.2% Triton X-100 for 10 minutes on ice. After permeabilization, cells were washed with PBS and blocked in 3%BSA, 0.1% Tween 20 in 4X saline sodium citrate buffer (SSC) for 1 hour at room temperature. Cells were then incubated with primary antibodies (**Table S4**) overnight at 4°C. Following PBS wash, cells were then incubated with Alexa-Fluor-488 or 568-conjugated secondary antibodies for 1 hour at room temperature, washed with PBS for 10 minutes 3 times, and stained with DAPI before mounting and imagining on LeicaDMI8 microscope at 100X magnification. ImageJ was used for image processing and quantification^75^. For assessment of micronuclei formation, DAPI staining was used to identify micronuclei-positive cells, which were quantified as a percentage of all cells analyzed.

### Western Blotting

Whole-cell lysates were prepared with RIPA buffer containing protease inhibitor (Sigma) and phosphatase inhibitor (Roche Applied Science) cocktail tablets and the protein concentration were determined by Bio-Rad Protein assay (Bio-Rad). Equivalent amounts of protein were resolved by SDS-PAGE and transferred to polyvinylidene fluoride microporous membrane (Millipore), blocked with 1.5% BSA in H20 containing 0.1% Tween-20 (TBS-T), and membranes were probed with the primary antibodies listed in (**Table S4**). Secondary antibodies were conjugated to horseradish peroxidase (HRP) and peroxidase activity was visualized using Chemiluminescent HRP substrate (Thermo Scientific). For quantification of protein samples, western blot images were analyzed using Image Lab Software for PC (v6.1). KEAP1 protein levels (Adj. Volume Int.) were normalized to GAPDH loading control and compared between samples.

### TCGA dataset analysis

mRNA expression levels were obtained from the TCGA cBioportal^76^ (RSEM - Batch normalized from Illumina HiSeq_RNASeqV2). Correlations of ARID1A and KEAP1 were analyzed using all samples (n = 10,071). For stratification by ARID1A-mutational status, ARID1A-mutated samples were sub-setted using VEG scores of “High” (meaning high impact on protein function - e.g., frameshift or loss). Cancer subtypes with at least 5 data points for either of ARID1A-WT or ARID1A-KO were selected for analysis.

### Differential gene expression analysis

RNA-seq data for RMG-1 cells isogenic for ARID1A was obtained from Wu *et. al*^22^. Differentially expressed genes were generated using DESEQ2^48^.

### Culture of patient-derived endometrial progenitor cells

Primary endometrial epithelial cells and organoids were derived as previously reported by Cochrane *et al.*^78^ from hysterectomy tissue from surgeries for non-cancer reasons performed at the Vancouver General Hospital (VGH) and University of British Columbia (UBC) hospitals. Tissues were collected as part of the OVCARE’s Gynaecologic Tissue Bank. These studies were approved by the Institutional Review Board (IRB) of UBC and British Columbia Cancer Agency (H05-60119), and use of the tissue for research purposes was approved by written informed consent by the patients.

Cells were seeded in 6 well plates to be transfected by ARID1A and non-targeting control CRISPR lentiviruses. ARID1A (12354, cell signaling, 1/1000) western blot was performed to confirm the downregulation of ARID1A protein in sgARID1A infected cells compared to NTC controls. Then, cells were seeded at a density of 1000 cells/well in 96 well plates. KEAP1 inhibitor treatment was performed after 24h using increasing concentrations and the plates were placed into an IncuCyte S3 live-cell imaging system contained in an incubator kept at 37 °C and 5% CO2 to assess proliferation for 7 days.

### Statistical analysis, data availability, and reproducibility

Statistical analysis was performed using GraphPad Prism 9. Experiments were repeated three times unless otherwise stated. The representative images were shown unless otherwise stated. Quantitative data were expressed as means ± SEM unless otherwise stated. Analysis of variance (ANOVA) was used to identify significant differences in multiple comparisons. Strains and plasmids are available upon request. The authors affirm that all data necessary for confirming the conclusions of the article are present within the article, figures, and tables.

## ACKNOWLEDGEMENTS

We thank DGH and Michael Anglesio for the RMG-1 ARID1A-WT and ARID1A-KO cells, as well as the other CCOC cell lines. We also thank Rodrigo Vallejos and Dawn Cochrane for their support with data analysis and organoid culture respectively. We also acknowledge Grace Cheng, Fraser Johnson, Dylan Farnsworth and the Morin Lab for their assistance with the IncuCyte. The pIND20-NRF2^E79Q^-HA construct was a generous gift from Bernard Weissman. Some of the results shown herein are in whole or part based upon data generated by the TCGA Research Network: https://www.cancer.gov/tcga. We also thank Arun Kumar for the insightful discussions that helped shape this manuscript. LAF would also like to acknowledge René Fournier, for providing continuous support through this work.

## FUNDING

LAF holds a CIHR Frederick Banting and Charles Best doctoral scholarship and a UBC Four Year Fellowship. This work was supported by a Terry Fox Research Institute Program Project grant to PCS, MH and DGH.

## DECLARATION OF INTERESTS

LAF is a member of the Office of the Canada Chief Science Advisor Youth Council (CSA-YC). PCS is CEO of Arrowsmith Genetics Inc.

## SUPPLEMENTAL MATERIAL LIST

***Supplemental Figure S1.* (A).** Representative western blot images of isogenic RMG-1 ARID1A-WT and KO cells. **(B-C)** Overlap of ARID1A-WT (**B**) or ARID1A-KO (**C**) fitness genes with the core essential gene training dataset (CEG2) for each replicate of our TKOv3 CRISPR screen. **(D)** Normalized read count data obtained using BAGEL2^72^ for all four TKOv3 sgRNAs targeting KEAP1 (n=2).

***Supplemental Figure S2.* (A)** Representative western blots showing KEAP1 siRNA knockdown efficiency in RMG-1 cells. **(B)** Severely impaired Growth of RMG-1 ARID1A-WT and KO cells treated with 100µM AI-1 as measured by IncuCyte S3 imaging system. Relative confluency presented as the growth from initial time point. Error bars represent SEM from triplicated wells from 3 independent experiments. *P* values obtained from extra sum-of-squares F test on calculated logistic growth rate are indicated on graph. **(C)** Representative western blot image of ARID1A, KEAP1 and NRF2 levels of the CCOC cell line panel. **(D)** Dose-response regression curves obtained from crystal violet proliferation assays for CCOC cell line panels. Confluency after 7 days of growth at indicated AI-1 concentrations was measured using crystal violet (averaged technical triplicate data from 3-4 independent experiments – or 2 independent experiments for 25µM -, mean ± SEM). IC50 values compared in **Figure 2C**. **(E)** *Upper left*: representative western blot images confirming ARID1A status in NTC or sgARID1A-transduced primary endometrium progenitor cells. *Right and below:* shows raw proliferation curves for individual patient derived cell cultures transduced with non-targeting (NTC5) or sgARID1A (ARID1A-KO) lentivirus and treated with AI-1 at indicated concentrations as measured by IncuCyte S3 imaging system (data presented as mean of quadruplicate wells from cells derived from a single patient). Note that despite different primary cell responses, the red ARID1A-KO curves are always more sensitive to AI-1.

***Supplemental Figure S3.* (A)** Quantification of KEAP1 mRNA levels (RSEM, log2) from pan-cancer TCGA data stratified by cancer subtype and ARID1A-status (significant p-values are displayed on graph). **(B)** Quantification of NRF2 and ARID1A mRNA levels (RSEM, log2) from pan-cancer TCGA data (n = 10,071) showing a mild positive correlation between the two genes. **(C)** Representative western blot image showing dose-dependent induction of NRF2^E79Q^ expression using doxycycline. **(D)** Representative images of crystal violet staining measuring proliferation of cells following 7 days of dox induction. **(E)** Quantification of RMG-1 ARID1A-WT and KO proliferation (normalized to DMSO control) showing no sensitivity to increasing concentrations of doxycycline in untransformed cells (mean ± SEM, n = 2 or 3 – see **Table S5**).**(F)** Representative western blot image showing NRF2 induction by AI-1 treatment but no effect on HDAC1 protein levels of RMG-1 cells treated with increasing concentrations of AI-1 for 24 hours.

***Supplemental Figure S4.* (A)** Representative western blot image showing increased γH2AX staining in sgARID1A-CRISPR cells after 48h treatment with AI-1 (data from cells derived from a single patient). **(B)** Quantification of confluency (relative to DMSO control) of RMG-1 cells treated with AI-1 (75µM)/ceralasertib (25nM) alone or in combination as measured after 60 hours of treatment by IncuCyte S3 imaging system (technical duplicate wells from three independent experiments; mean ± SEM, ANOVA, significant p-values displayed on graph).

***Supplemental Figure S5.*** Raw Western blot images.

***Supplemental Table 1 – CRISPR screen datasets***

***Sheet 1:*** Normalized read counts from the analysis of biological replicate 1 of our TKOv3 CRISPR screen. ***Sheet 2:*** Normalized read counts from the analysis of biological replicate 2 of our TKOv3 CRISPR screen. ***Sheet 3:*** Fold change of sgRNAs from the analysis of biological replicate 1 of our TKOv3 CRISPR screen. ***Sheet 4:*** Fold change of sgRNAs from the analysis of biological replicate 2 of our TKOv3 CRISPR screen. ***Sheet 5:*** BAGEL output of Bayes Factor, Precision, Recall and FDR values of fitness genes in ARID1A-WT and KO cells (replicate 1). ***Sheet 6:*** BAGEL output of Bayes Factor, Precision, Recall and FDR values of fitness genes in ARID1A-WT and KO cells (replicate 2). ***Sheet 7:*** Summary of all significantly enriched fitness genes identified in ARID1A-KO cells. ***Sheet 8:*** ARID1A-KO-specific hits for each biological replicate of the TKOv3 screen (i.e. synthetic lethal partners of ARID1A), along with overlap ARID1A-SL consensus from the two replicates.

***Supplemental Table 2 – GO analysis (David v6.8)***

***Sheet 1:*** GO analysis of enriched biological processes for the ARID1A fitness genes identified in both CRISPR screen replicates (overlap presented in **Supplemental Table 1.7**). ***Sheet 2:*** GO analysis of enriched biological processes for the consensus ARID1A-SL dataset (103 genes, **Supplemental Table 1.8**).

***Supplemental Table 3 -Differential gene expression***

Differential gene expression analysis of up-regulated genes (***Sheet 1***) and down-regulated genes (***Sheet 2***) in RMG-1 ARID1A-KO cells from Wu *et al.*^22^ using DESEQ2^47^.

***Supplemental Table 4 - Miscellaneous***

***Sheet 1:*** Reagent List. ***Sheet 2***: p.IND20-NRF2-E79Q-HA plasmid sequence.

***Supplemental Table 5 - Raw Data***

Raw data for each Figure presented in separate sheets.

## Notes

### Summary of Updates

The RPE1 cell line data was removed. The figures were revised to better represent variability in the primary cell data. New analysis was added to Figure 3.

