## Supplemental Figures and Tables for "Genome-wide CRISPR screen identifies KEAP1 perturbation as a vulnerability of ARID1A-deficient cells": Supplemental Figure 5_final (raw blot images).pdf

Original Blots

### Figure 3E

ARID1A

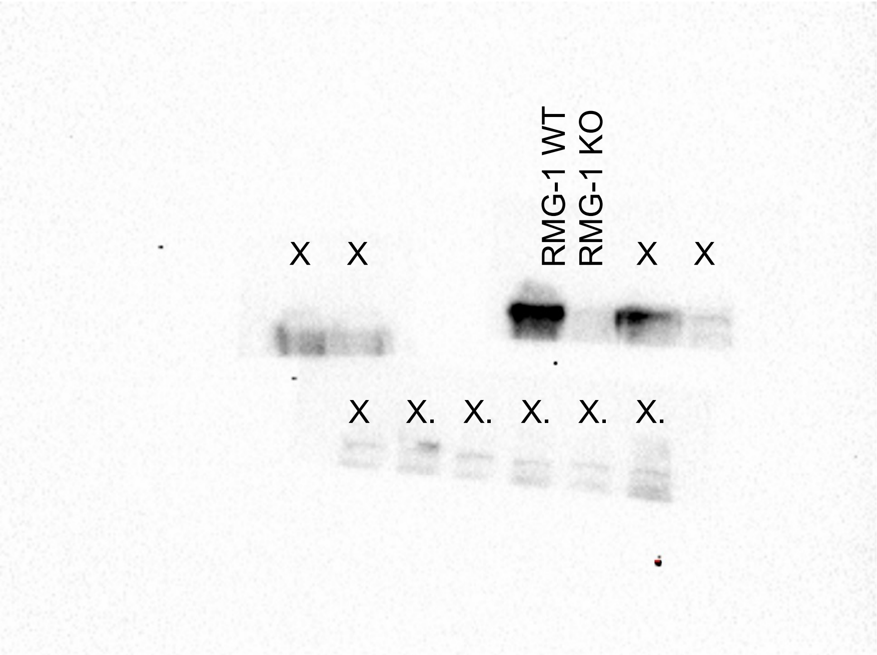


KEAP1

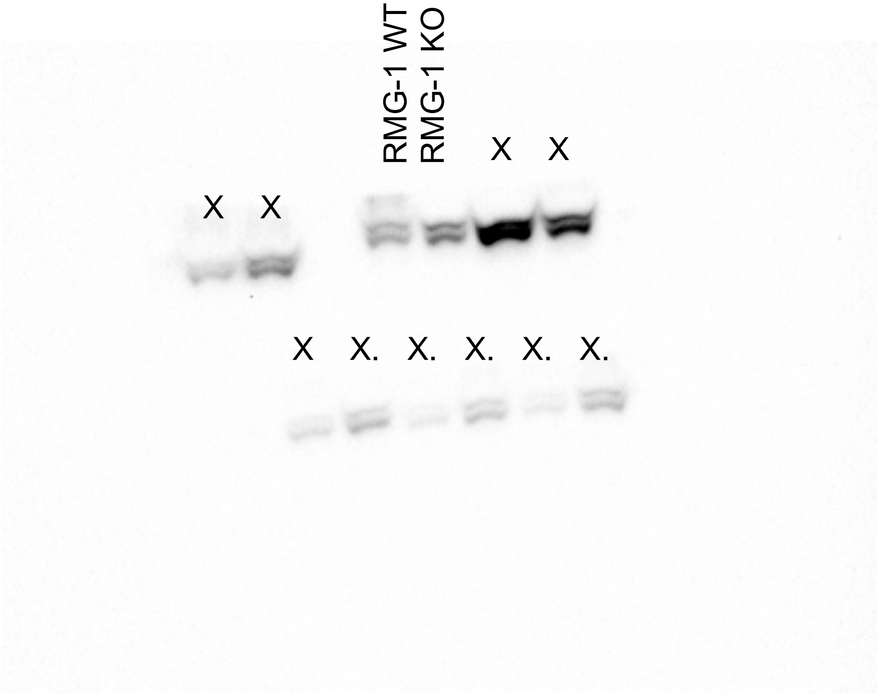


GAPDH

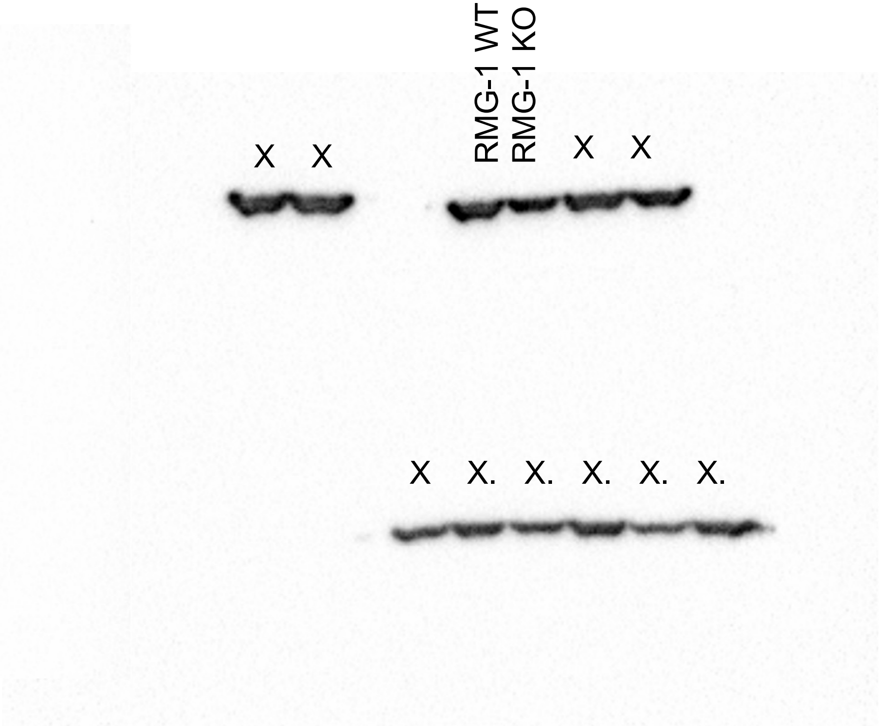


X = not the gel/sample of interest.

### Figure S1A

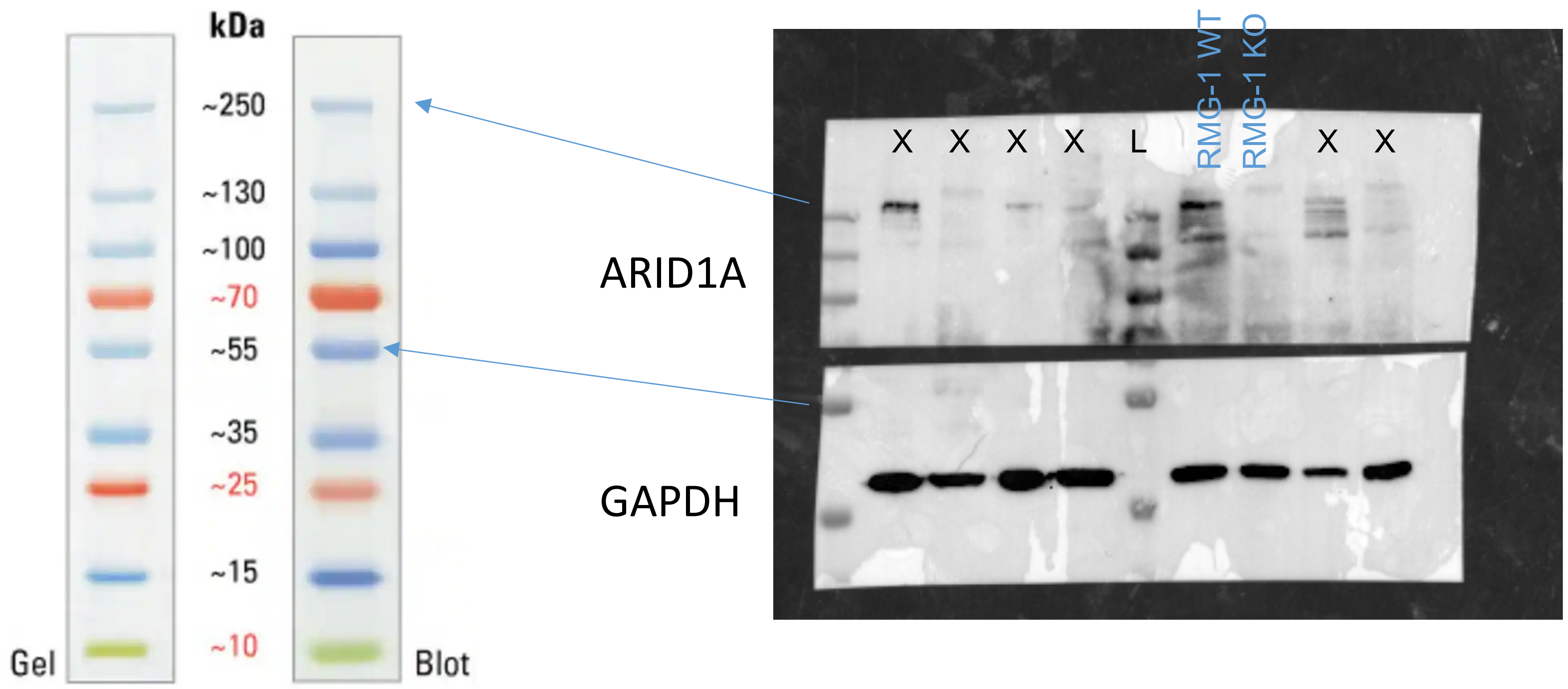


X = not the gel/sample of interest.

### Figure S2A

KEAP1 & NRF2

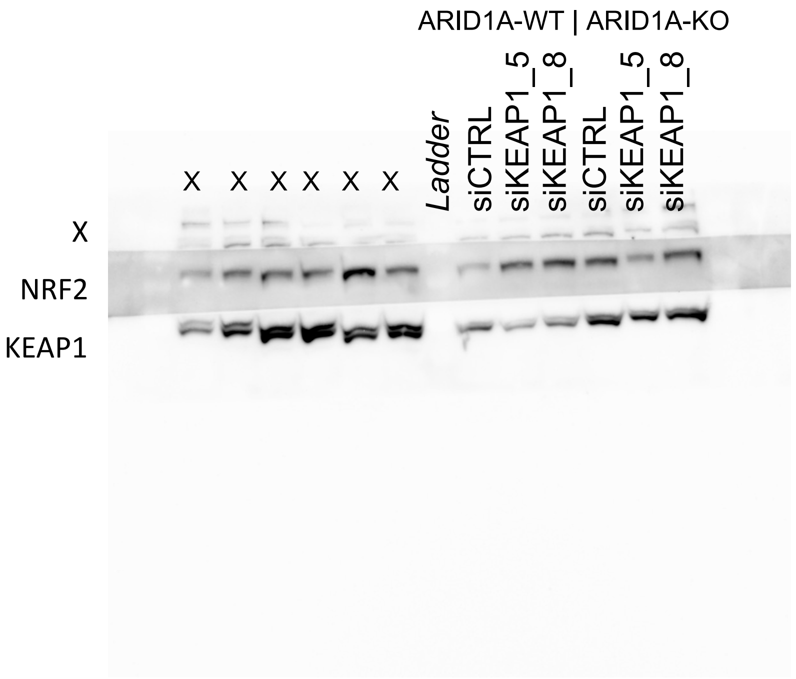


GAPDH

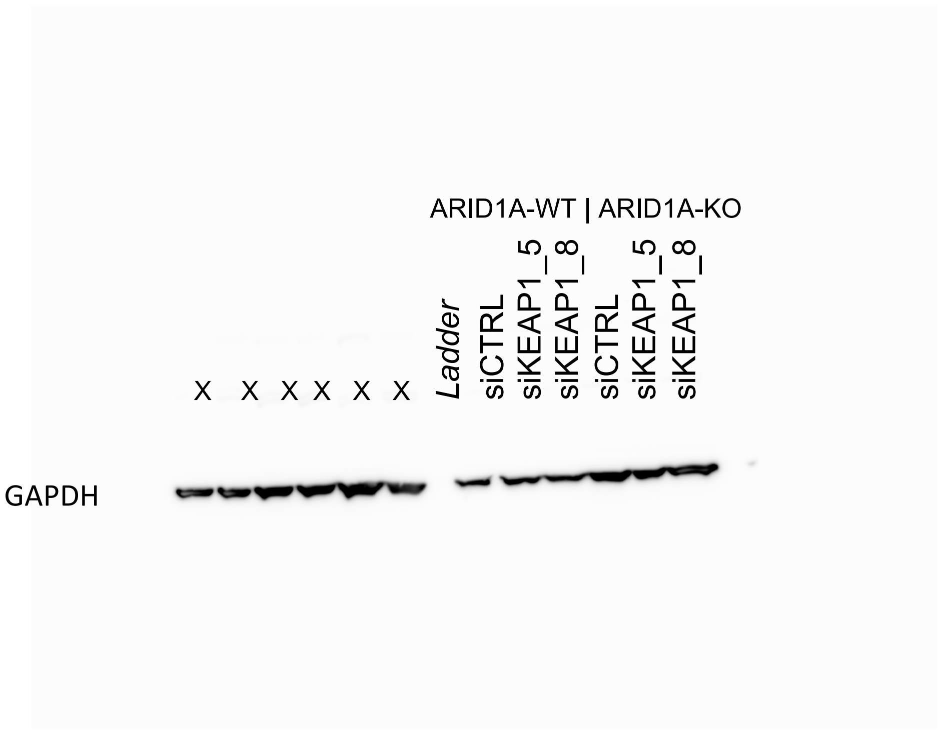


X = not the gel/sample of interest.

### Figure S2C

ARID1A

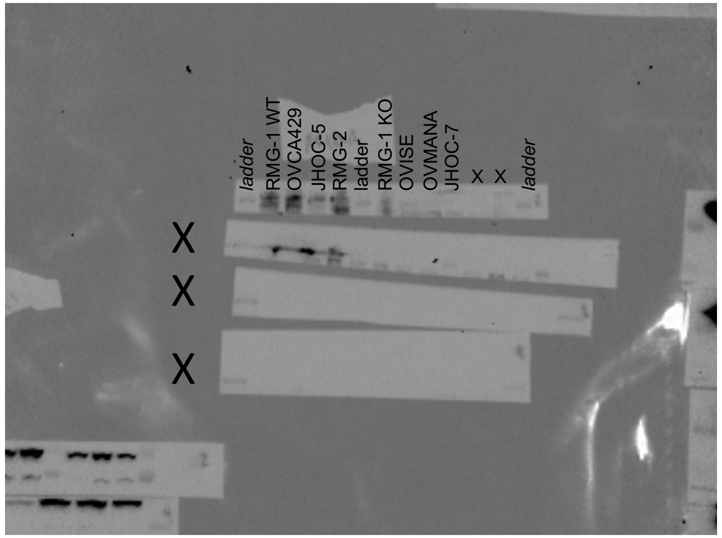


NRF2

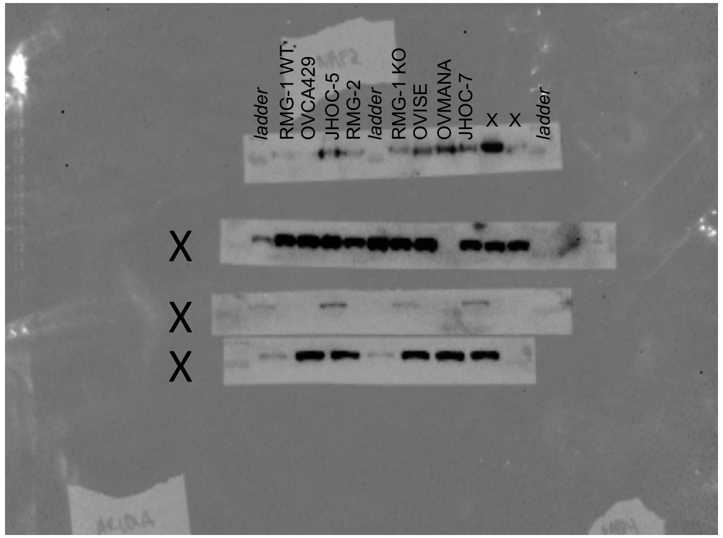


KEAP1

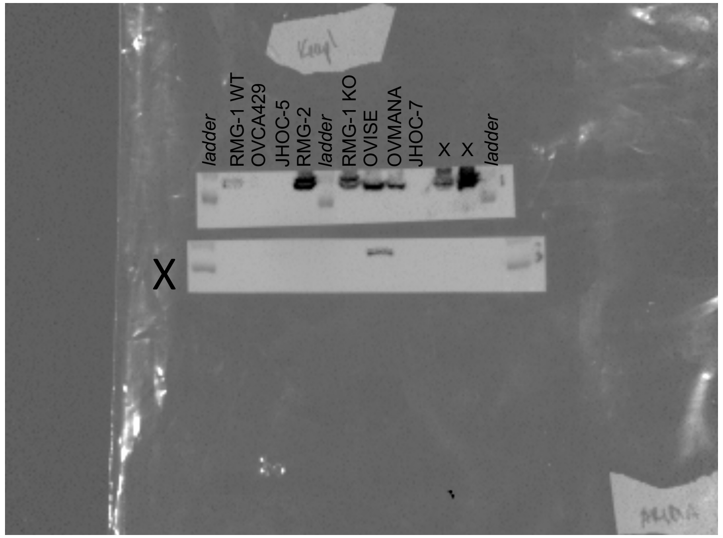


GAPDH

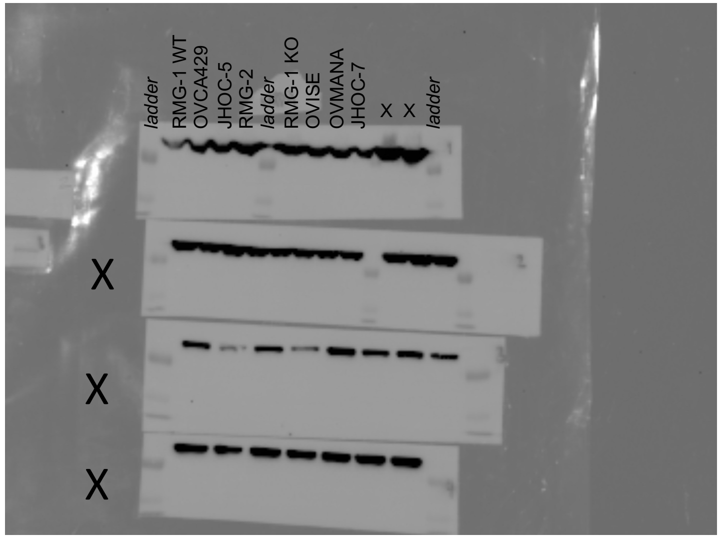


X = not the gel/sample of interest.

### Figure S2E

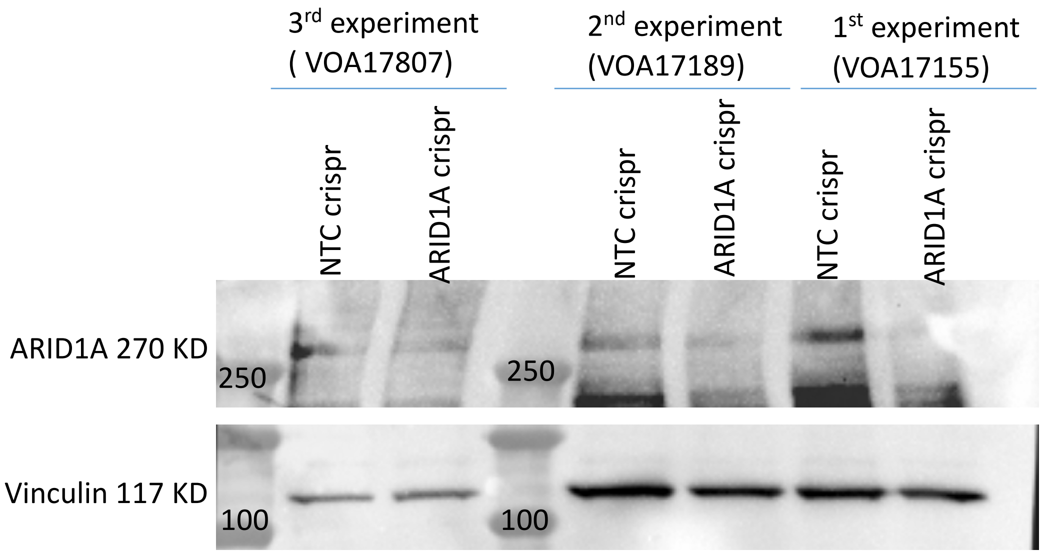
3rd experiment
( VOA17807)
2nd experiment
(VOA17189)
1st experiment
(VOA17155)
NTC crispr
ARID1A crispr
NTC crispr
ARID1A crispr
NTC crispr
ARID1A crispr
ARID1A 270 KD
250
250
Vinculin 117 KD
100
100

### Figure S3C

HA

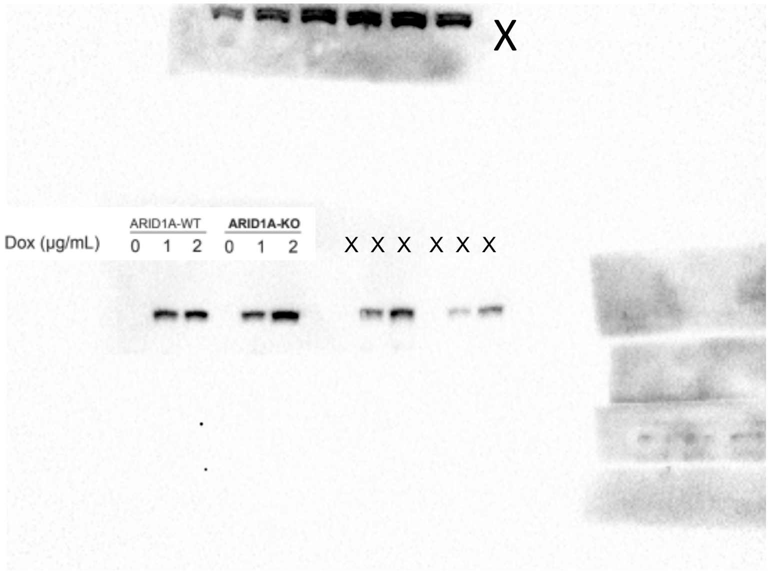


KEAP1

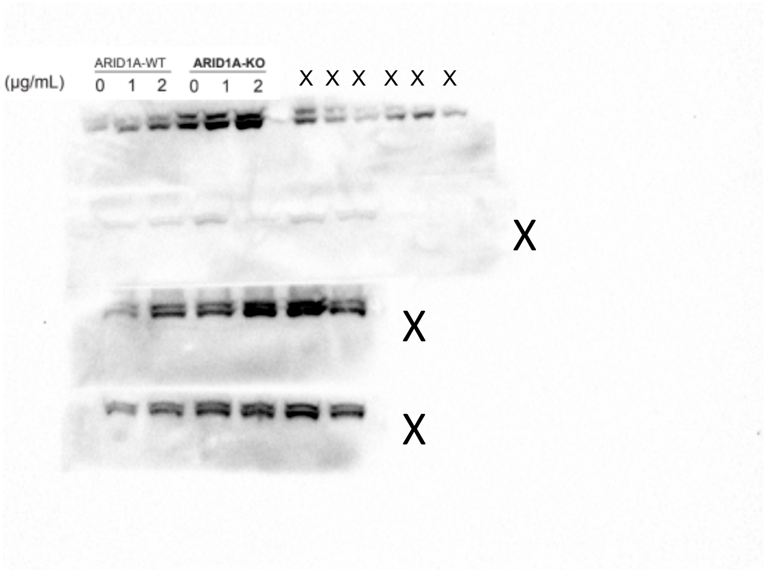


GAPDH

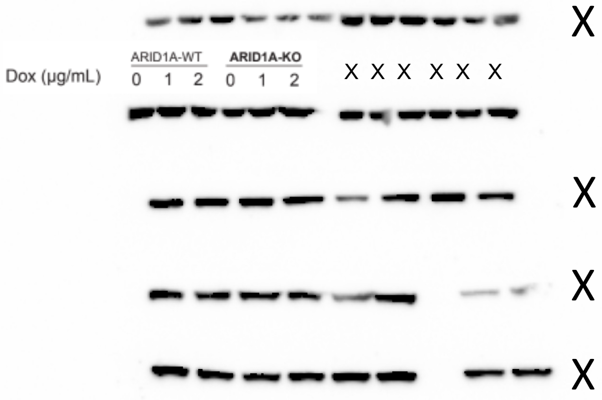


X = not the gel/sample of interest.

### Figure S3F

NRF2 HDAC1 GAPDH

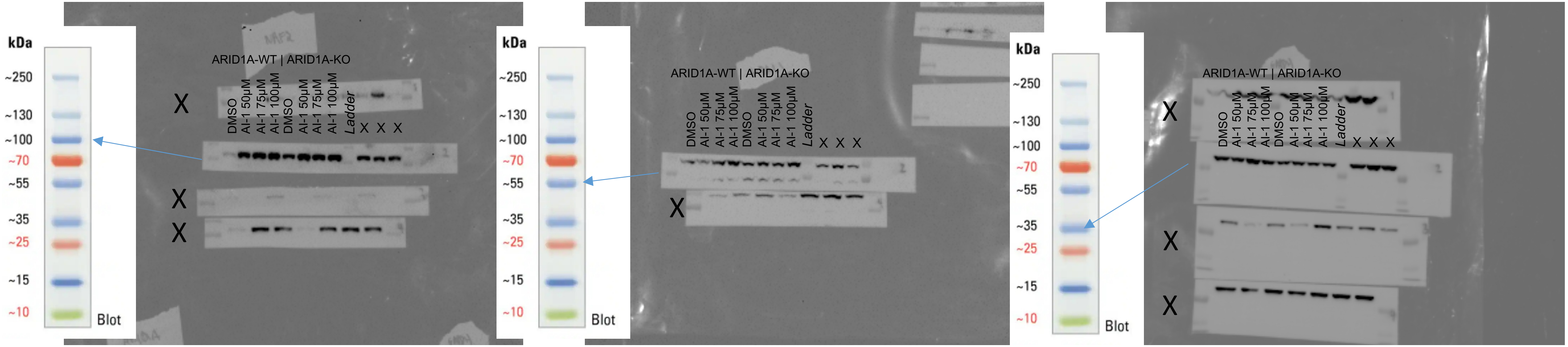


X = not the gel/sample of interest.

### Figure S4A

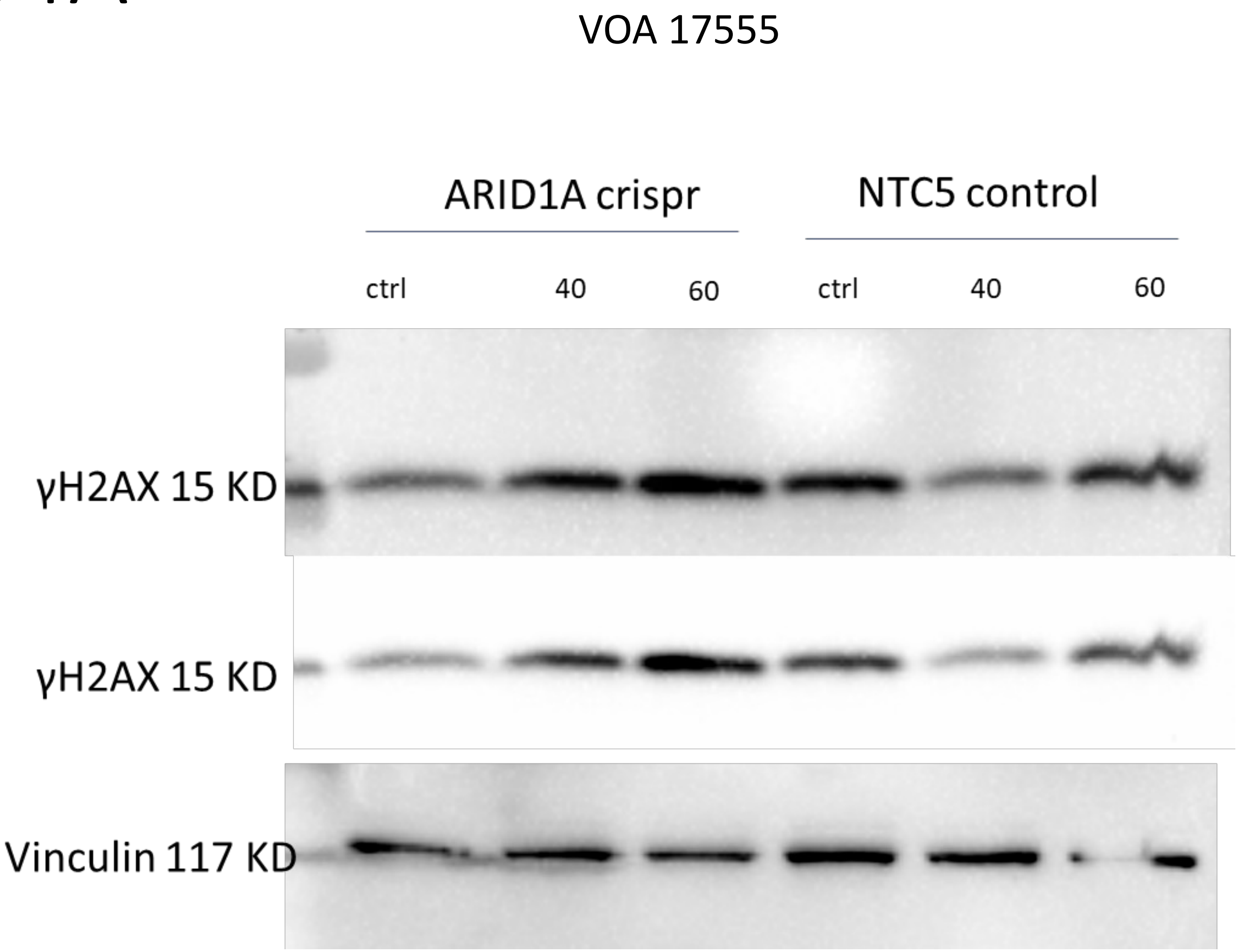
VOA 17555
ARID1A crispr
NTC5 control
ctrl
40
60
ctrl
40
60
γH2AX 15 KD
γH2AX 15 KD
Vinculin 117 KD

### Appendix

Additional images

### Figure S2C

ARID1A

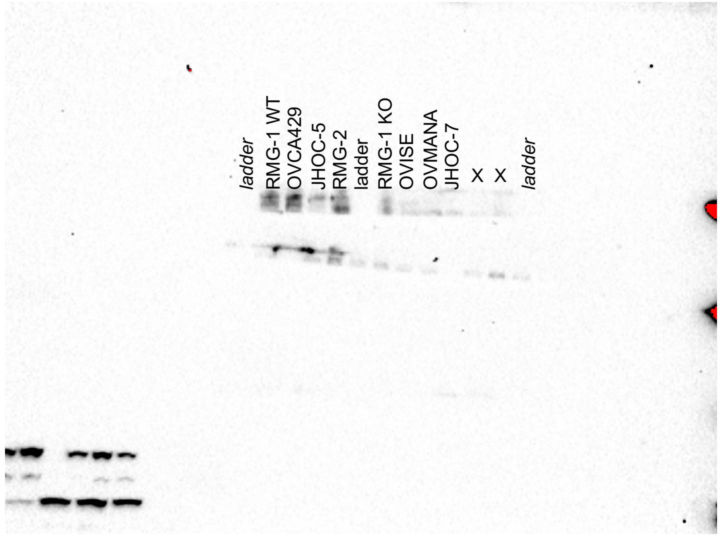


NRF2

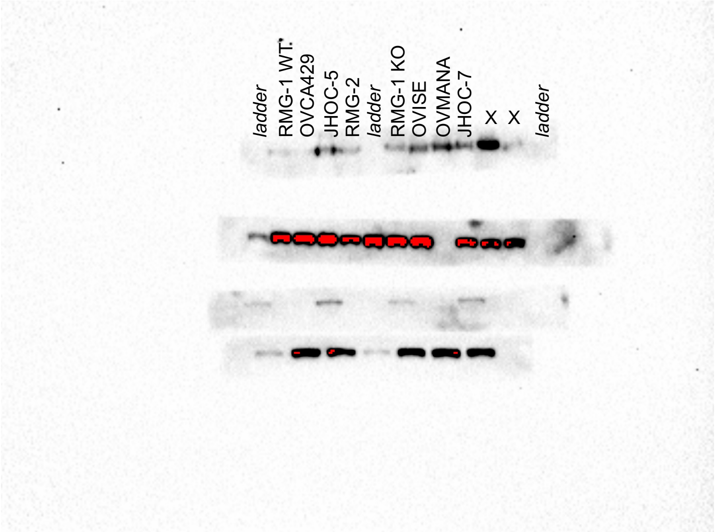


KEAP1

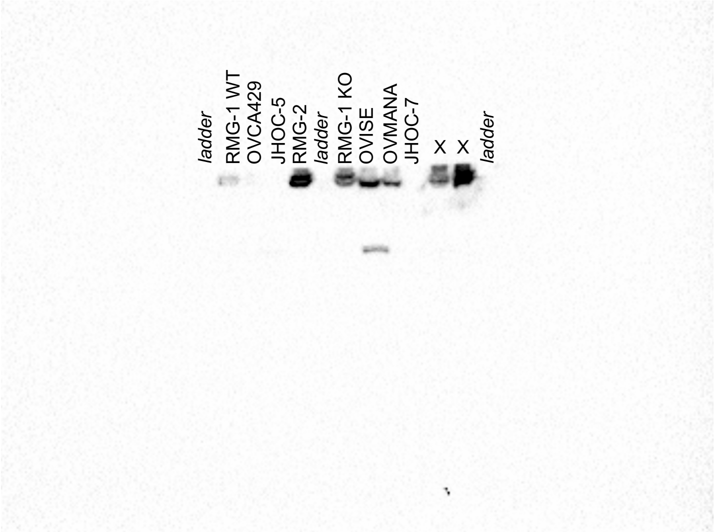


GAPDH

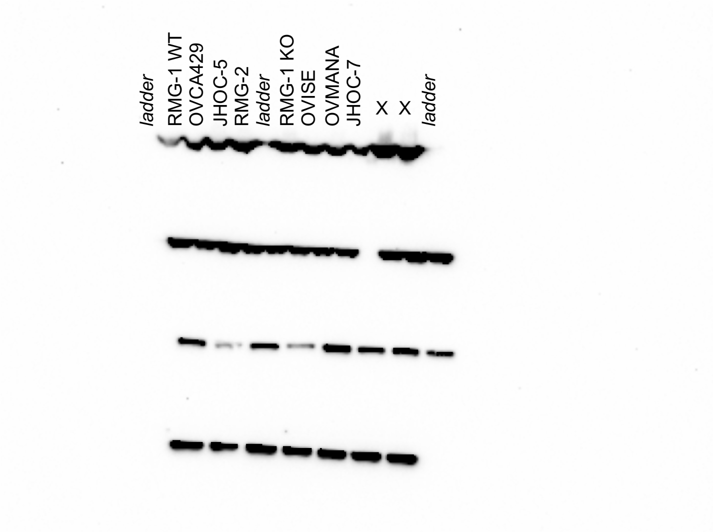


X = not the gel/sample of interest.

### Figure S3F

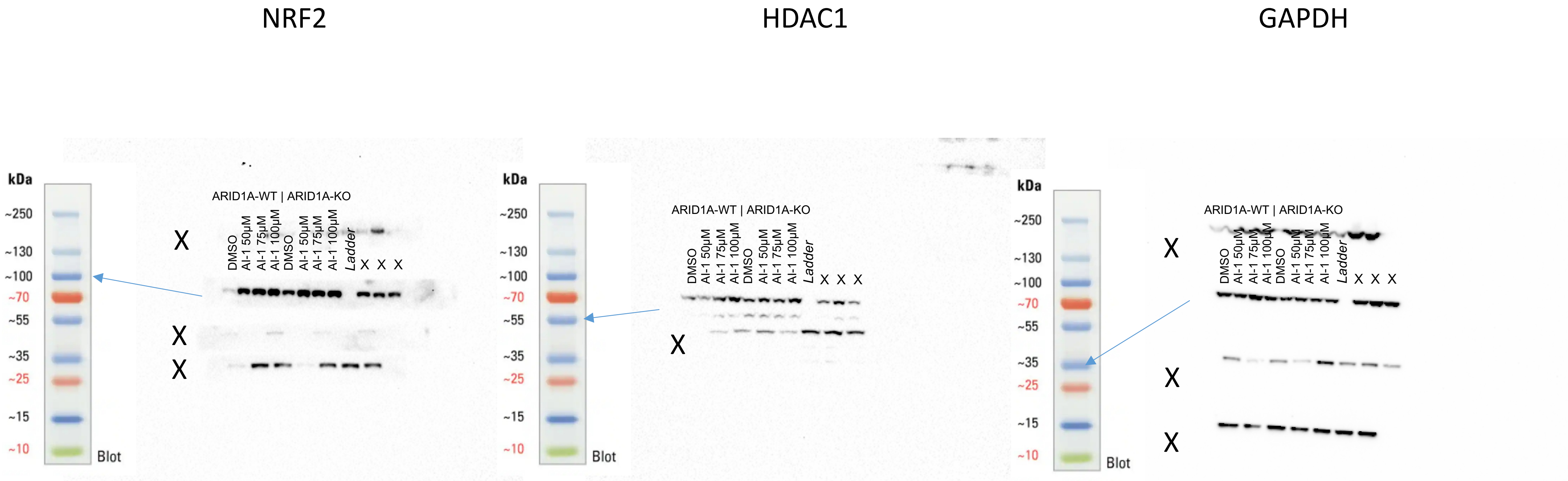


X = not the gel/sample of interest.
