## Supplementary figures and images for "Genome-wide CRISPR screen identifies KEAP1 perturbation as a vulnerability of ARID1A-deficient cells"

### Supplemental Figure 1_Final.pdf

Supplemental Figure 1

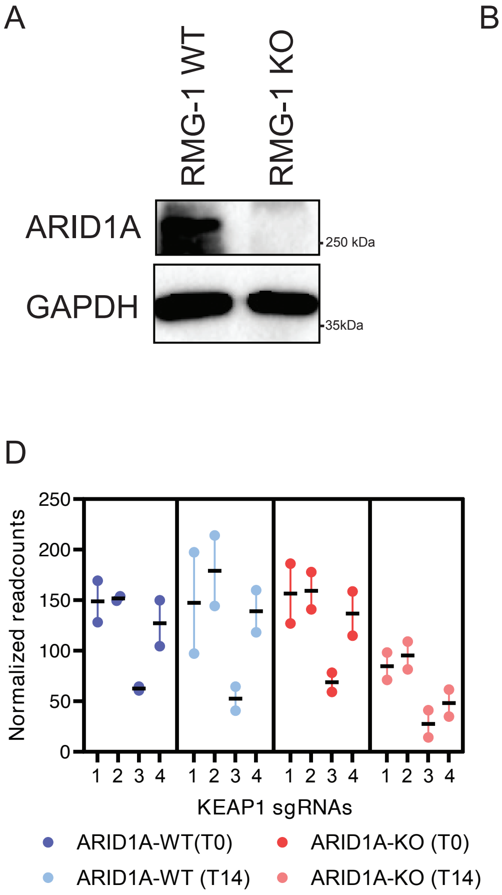

### Supplemental Figure 2_final.pdf

#### Supplemental Figure 2

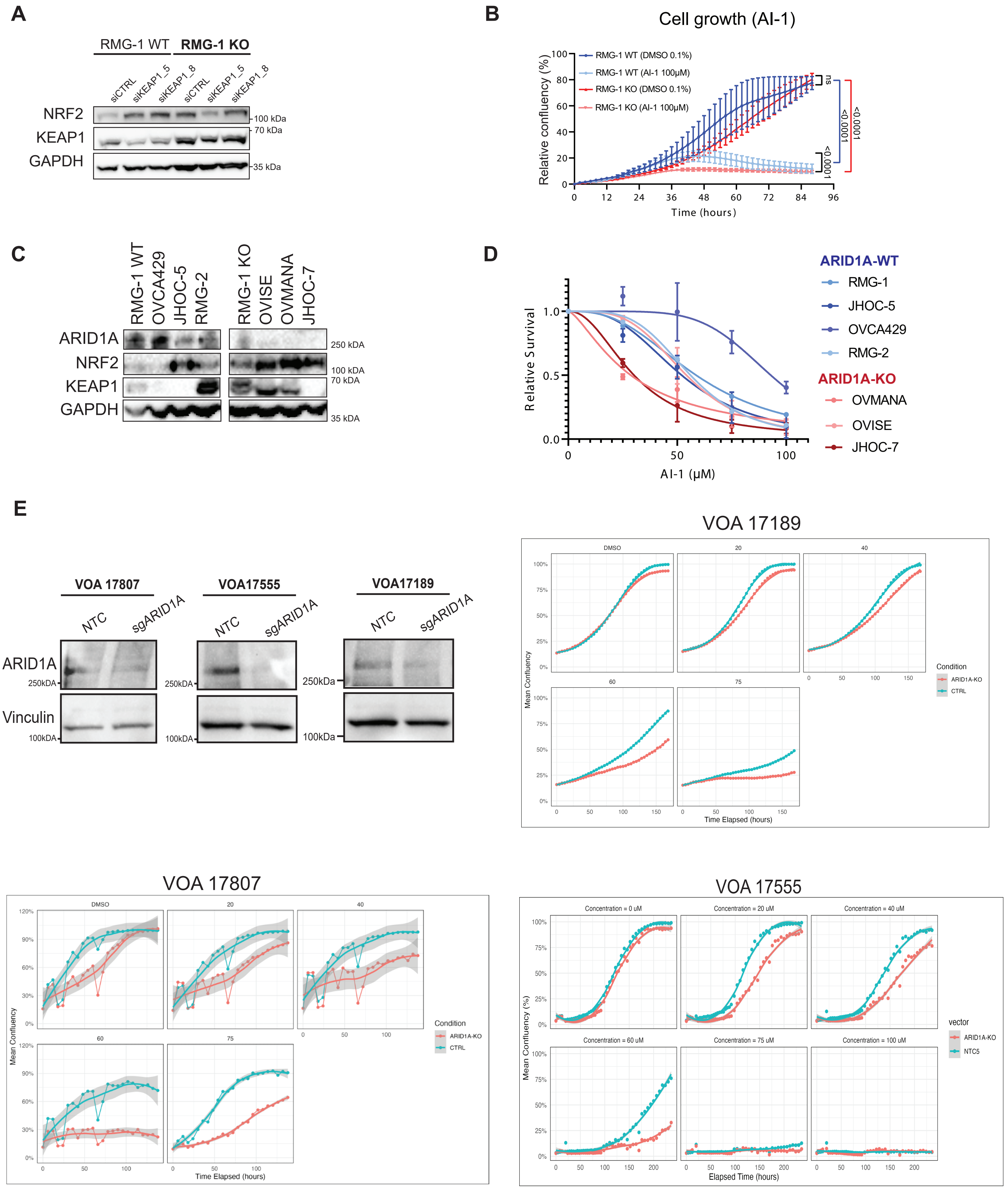

### Supplemental Figure 3_final.pdf

Supplemental Figure 3

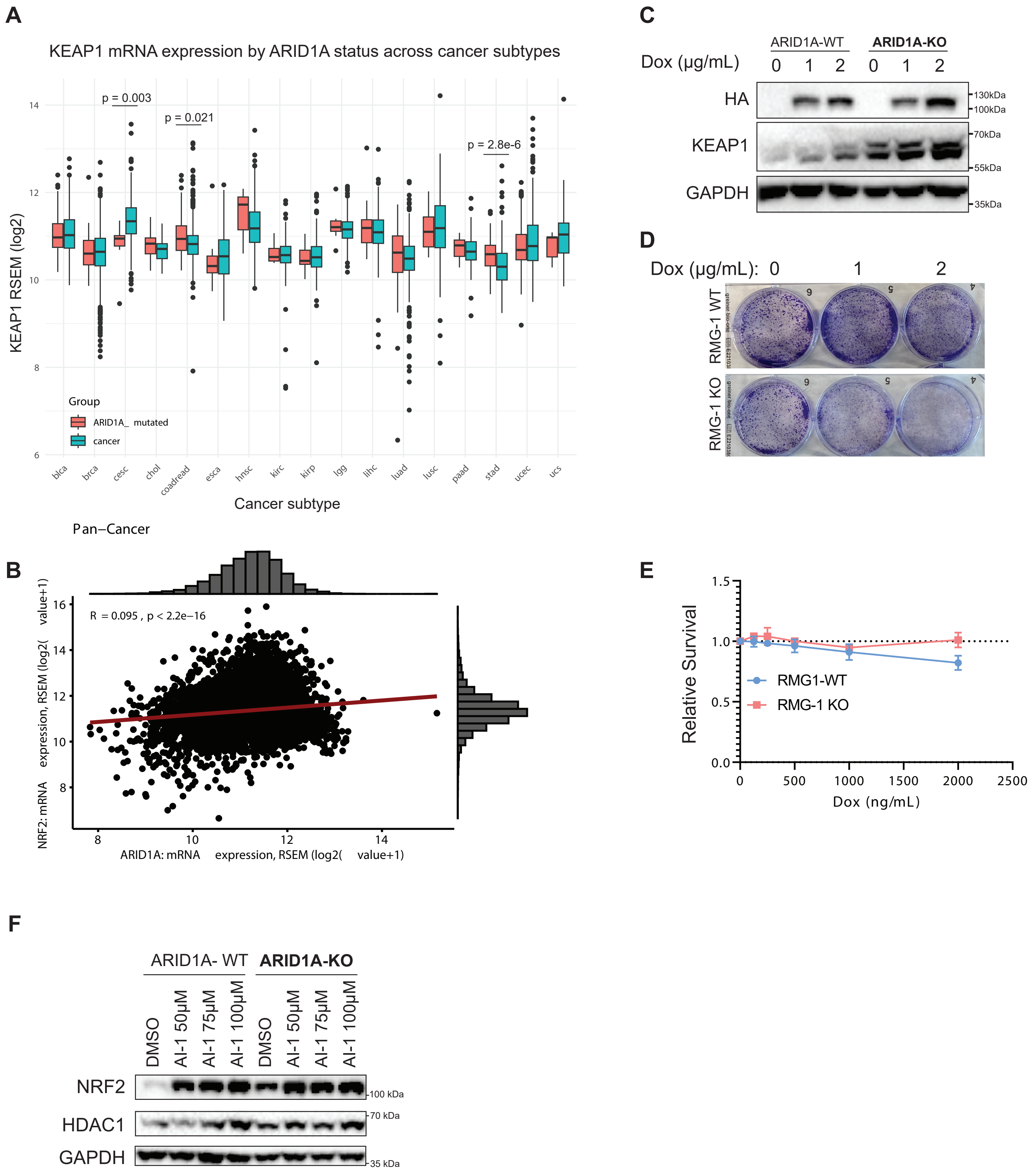

### Supplemental Figure 4_final.pdf

Supplemental Figure 4

A

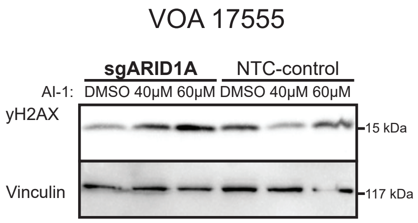


B

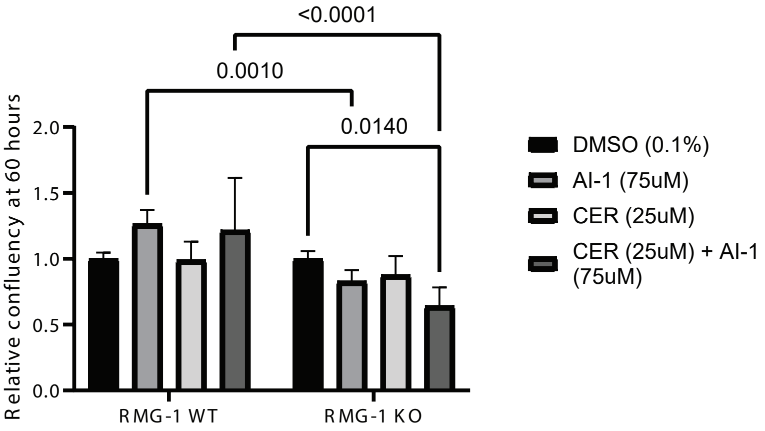
